# Targeting the dependence on PIK3C3-mTORC1 signaling in dormancy-prone breast cancer cells blunts metastasis initiation

**DOI:** 10.1101/2023.08.02.551681

**Authors:** Islam E. Elkholi, Amélie Robert, Camille Malouf, Hellen Kuasne, Stanislav Drapela, Graham Macleod, Steven Hébert, Alain Pacis, Virginie Calderon, Claudia L. Kleinman, Ana P. Gomes, Julio A. Aguirre-Ghiso, Morag Park, Stéphane Angers, Jean-François Côté

**Affiliations:** Montreal Clinical Research Institute, Montreal, Canada; Molecular Biology Programs, Université de Montréal, Canada; Rosalind and Morris Goodman Cancer Institute, McGill University, Canada; Department of Molecular Oncology, H. Lee Moffitt Cancer Center & Research Institute, Tampa, USA; Donnelly Centre for Cellular & Biomolecular Research, University of Toronto, Canada; Leslie Dan Faculty of Pharmacy, University of Toronto, Canada; Lady Davis Institute for Medical Research, Jewish General Hospital, Montreal, Canada; Canadian Centre for Computational Genomics, McGill University, Montreal, Canada; Department of Human Genetics, McGill University, Canada; Department of Cell Biology, Albert Einstein College of Medicine, Bronx, USA; Cancer Dormancy and Tumor Microenvironment Institute, Albert Einstein Cancer Center, Albert Einstein College of Medicine, Bronx, USA; Ruth L. & David S. Gottesman Institute for Stem Cell Research & Regenerative Medicine, Albert Einstein Cancer Center, Albert Einstein College of Medicine, Bronx, USA; Institute for Aging Research, Albert Einstein Cancer Center, Albert Einstein College of Medicine, Bronx, USA; Department of Biochemistry, Temerty Faculty of Medicine, University of Toronto, Canada; Department of Biochemistry and Molecular Medicine, Université de Montréal, Canada; Department of Anatomy and Cell Biology, McGill University, Canada

## Abstract

Halting breast cancer metastatic relapses following primary tumor removal and the clinical dormant phase, remains challenging, due to a lack of specific vulnerabilities to target during dormancy. To address this, we conducted genome-wide CRISPR screens on two breast cancer cell lines with distinct dormancy properties: 4T1 (short-term dormancy) and 4T07 (prolonged dormancy). We discovered that loss of class-III PI3K, Pik3c3, revealed a unique vulnerability in 4T07 cells. Surprisingly, dormancy-prone 4T07 cells exhibited higher mTORC1 activity than 4T1 cells, due to lysosome-dependent signaling occurring at the cell periphery. Pharmacological inhibition of Pik3c3 counteracted this phenotype in 4T07 cells, and selectively reduced metastasis burden only in the 4T07 dormancy-prone model. This mechanism was also detected in human breast cancer cell lines in addition to a breast cancer patient-derived xenograft supporting that it may be relevant in humans. Our findings suggest dormant cancer cell-initiated metastasis may be prevented in patients carrying tumor cells that display PIK3C3-peripheral lysosomal signaling to mTORC1.

**Statement of Significance:** We reveal that dormancy-prone breast cancer cells depend on the class III PI3K to mediate a constant peripheral lysosomal positioning and mTORC1 hyperactivity. Targeting this pathway might blunt breast cancer metastasis.

## INTRODUCTION

Nearly one-quarter of breast cancer (BC) patients experience metastatic relapse, months to over two decades after their initial diagnosis due to metastatic dormancy (1–3). The latter phenomenon occurs when tumor cells in secondary organs become quiescencent (cellular dormancy) or maintain a balance between proliferation and apoptosis (micrometasatatic dormancy), leading to clinically undetectable metastatic disease. The understanding of the molecular and biological dynamics of metastatic dormancy is still emerging (4). Identifying the molecular determinants that mediate the survival mechanisms of dormant disseminated tumor cells (DTCs) and their transition to metastatic outgrowth is needed to develop therapies to prevent relapse. These molecular determinants include cancer cell intrinsic factors and microenvironmental cues that can direct the fate and response of DTCs to therapies (5–8).

Different human and murine BC cell line models with shared origins but differential metastatic behaviors have been invaluable in indentifying factors that mediate DTCs’ fate (reviewed in (8)). One example is the 4T1 and 4T07 murine BC cell lines, originating from the same spontaneous metastatic nodule in BALB/c mice (9). In immunocompetent BALB/c mice, spontaneously disseminated 4T1 cells form overt lung metastases after a short latency period, while 4T07 cells remain dormant and show low frequency of metastasis. In immunocompromised mice, disseminated 4T07 cells escape dormancy and form overt metastases, though less aggressively than 4T1 cells (10), indicating intrinsic differences between the two cell lines. Indeed, differential gene expression analysis between 4T1 and 4T07 tumors identified TWIST1 as an upregulated gene in 4T1 cells mediating epithelial-mesenchymal transition and metastasis (11). Moreover, a gain-of-function screen using a cDNA library derived from 4T1 identified Coco as a mediator of reactivation of disseminated dormant 4T07 cells (12). The parallel study of these two cell lines therefore offers an opportunity to define new cancer cell intrinsic factors that contribute to metastatic progression.

In this study, we investigated whether the 4T1 and 4T07 cells exhibit differential activity in signaling pathways that could be exploited for therapeutic intervention *in vivo*. Using a genome wide loss-of-function CRISPR screening framework, we systematically identified the fitness genes that are differentially essential for the growth of 4T1 versus 4T07 cells. Identifying fitness genes involved in a specific signaling pathway indicates potential activity of this pathway and sensitivity to its targeted inhibition (13). We discovered that 4T1, but not 4T07, cells rely on and show higher activity of class I PI3K. Interestingly, 4T07 cells demonstrate higher activity of the downstream mechanistic Target of Rapamycin complex 1 (mTORC1). We show that 4T07, unlike 4T1 cells, depend on class III PI3K to maintain peripheral lysosomal positioning, which is linked to mTORC1 hyperactivity. This lysosome positioning-associated PIK3C3-mTORC1 signaling circuit is present in human BC cell lines, including a patient-derived xenograft line originating from a patient with BC metastatic relapse. Notably, selective inhibition of Pik3c3 exclusively kills the dissiminated 4T07 cells. Our findings suggest that targeting the PIK3C3-mTORC1 signaling axis could be a promising therapeutic strategy to reduce metastatic burden in some BC patients by eliminating DTCs.

## MATERIALS AND METHODS

### Cell lines, PDX line, and reagents

Parental 4T07 cells (9) were obtained from the Karmanos Cancer Institute, Wayne State University. The 4T07-TGL cells (12) (Memorial Sloan Kettering Cancer Center, MSKCC) were obtained from Dr. Filippo Giancotti (Herbert Irving Comprehensive Cancer Center, HICCC). 4T1 cells were obtained from ATCC. To generate 4T1-TGL cells, HEK293T cells were seeded in 6 cm plates and transfected with 3 ug of TGL vector (14) (gift from Dr. Vladimir Ponomarev, MSKCC) and 3 ug of pCL-Ampho using CaCl_2_ towards retroviral production. Then parental 4T1 cells were infected and stable 4T1-TGL cells were generated and established after sorting for GFP positive cells. To generate 4T07 cells expressing doxycycline-inducible shRNA, we first generated pLKO-Tet-ON plasmids encoding different sequences predicted to either target *Pik3c3* or act as a non-targeting control (see table below for sequences). HEK293FT cells were then transfected with the generated plasmids in addition to psPAX2, pMD2.G to produce lentiviruses. 4T07 cells were then transduced with these lentiviruses and selected with puromycin (3 µg/ml) for 3-5 days. 4T07-shRNA cell lines were induced with doxycycline (1 µg/ml). All the 4T1 and 4T07 cell lines in addition to T47D cells were cultured in Dulbecco’s modified Eagle’s medium (DMEM; Wisent #319-005-CL), supplemented with 10% fetal bovine serum (FBS; Gibco #12483-020), and 1% penicillin/streptomycin (P/S) antibiotics (Wisent #450-201-EL). CAMA-1 and HCC70 were cultured in Eagle’s minimum essential medium (EMEM; Wisent #320-026-CL) and Roswell Park Memorial Institute 1640 (RPMI; Wisent #350-000-CL) supplemented with 10% FBS and 1% P/S.

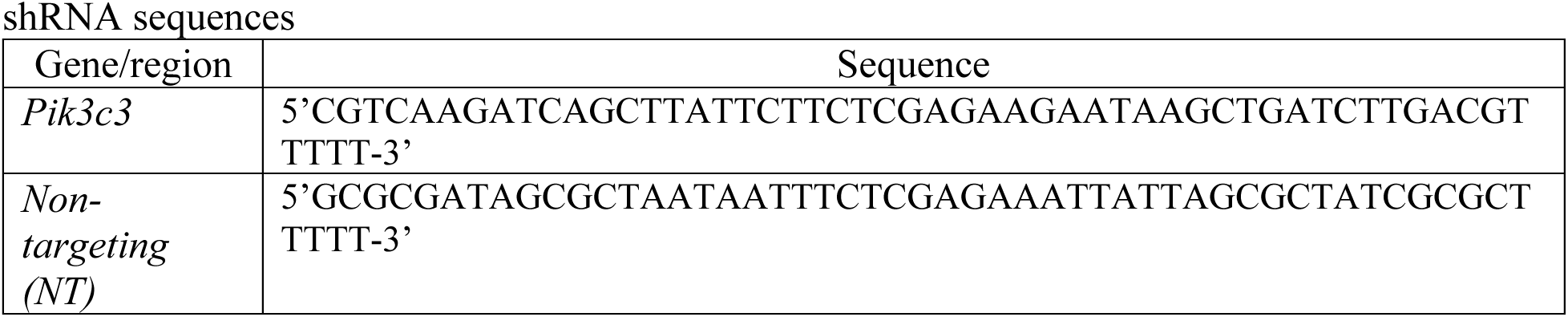

For serum starvation, cells were cultured overnight (O/N) in serum-free DMEM supplemented with 1% P/S. For amino acids starvation, cells were first serum starved O/N, then rinsed with PBS and incubated in Earle’s Balanced Salt Solution (EBSS; ThermoFisher Scientific #24010043)) for two hours at 37 °C. Refeeding with serum was done by replacing the serum-free DMEM with 10% FBS-supplied DMEM. To refeed with insulin, 100 nM insulin (Humulin R U-100) was added to the serum-free DMEM, and cells were incubated for 30 minutes at 37 °C. To refeed with amino acids, MEM Amino Acids solution (50x; Sigma-Aldrich #M5550) was added to the EBSS (1:50), and cells were incubated for 30 minutes at 37 °C.

All patient-derived xenograft (PDX) epithelial cell lines were derived from tissues obtained according to the ethical regulations of McGill University and after informed consents, as stated previously (15). PDX 1915 (15) was cultured in DMEM (Gibco #11995-065) and F-12 Nutrient Mixture (Ham) (Gibco #11765-054) (1:4), 5% FBS (Life Technologies, #12483012), 0.4 µg/mL Hydrocortisone (Sigma-Aldrich #H0888-1G), 5 µg/mL Insulin (Gibco #12585-014), 8.4 ng/mL Cholera toxin (Sigma-Aldrich #C8052), 10 ng/mL Epidermal growth factor (BPS bioscience #90201-1), 10 µM Y-27632 (Abmole #M1817), 50 µg/mL Gentamicin (Gibco #15710-072), 1% P/S (Thermo Fisher Scientific, #15140-122), Amphotericin B (1 µg/ml) (Thermo Fisher Scientific, #15290026).

The PIK3CA-inhibitor BYL719 (ChemieTek #CT-BYL719), AKT-inhibitor MK-2206 2HCl (Selleck Chemicals #S1078), and VPS34-IN1 (Cayman Chemical #17392) were dissolved in DMSO and used in different concentrations as indicated.

*Mycoplasma testing.* Parental 4T07 cells were mycoplasma-negative as last tested by PCR (see primers sequences below) in October 2022. Given that 4T07-TGL cells were contaminated with mycoplasma upon receiving them, they were treated with Plasmoscin (InvivoGen #ant-mpt-1) for two weeks as per the manufacturer’s protocol before using them in experiments described in this work. 4T1 cells were mycoplasma-negative as last tested by PCR in October 2022. PDX-1915 was tested for mycoplasma, before being used for experiments, using the Mycoalert detection kit (Lonza #LT07-318) as per the manufcaturer’s instructions. T47D, CAMA-1, and HCC70 cells were not tested for mycoplasma contamination by PCR. However, it should be noted that all the described cell lines were frequently used in immunofluroscence experiments where DAPI staining was performed and verified mycoplasma-negative status for all the lines included in this work. For PCR reaction, we used a mixture of six forward primers (CGCCTGAGTAGTACGTTCGC; CGCCTGAGTAGTACGTACGC; TGCCTGAGTAGTACATTCGC; TGCCTGGGTAGTACATTCGC; CGCCTGGGTAGTACATTCGC; CGCCTGAGTAGTATGCTCGC) and a mixture of three reverse primers (GCGGTGTGTACAAGACCCGA; GCGGTGTGTACAAAACCCGA; GCGGTGTGTACAAACCCCGA). In all the conducted tests, a positive (mycoplasma-positive sample) and a negative control were used to verify the test. This protocol was adapted from (https://bitesizebio.com/23682/homemade-pcr-test-for-mycoplasma-contamination/) based on these references (16,17).

### CRISPR-Cas9 Screening

The mouse Genome-Scale CRISPR Knock-Out (mGeCKO) library A (Addgene #1000000052), composed of ∼68,000 gRNAs: 3 gRNAs/gene, 4 gRNAs/miRNA, and 1000 nontargeting sequence, was amplified and prepared for NGS (MiSeq Illumina) to assess the gRNA distribution and library quality, as previously described (18). In reference to the original library, the assessed parameters of the amplified library (percentage of undetected gRNAs and perfectly matching gRNAs in addition to the skew ratio of the top 10% represented gRNAs to bottom 10%) were all within the recommended ranges for a reliable library (18).

Lentiviral production was performed as previously described (18). Briefly, low passage HEK 293FT cells were plated in Nunc EasYFlask cell culture flasks (Thermo Fisher Scientific # 156340) and transfected, using lipofectamine 2000 (Thermo Fisher Scientific #<u>11668027</u>), with psPAX2 (23.4 µg), pMD2.G (15.3 µg), and 30.6 µg of the lentiCRISPR V2 vector encompassing the Cas9 in addition to the amplified library. 48 hours later, the viruses-containing media was collected and filtered using 0.45 um sterile filters (Sarstedt #83.1826). A multiplicity of infection (MOI) test was performed to define the optimum volume of the viruses-containing media to be used for the screens.

The 4T1 and 4T07 cells were infected independently, as previously described (18). Briefly, the two cell lines were seeded in 12 well plates (1.5-2 x 10^6^ cells/well) and then subjected to spinfection (MOI < 0.3) by centrifugation at 2,000 RPM for 2 hours at 33°C in the presence of polybrene (10 µg/ml) (Sigma-Aldrich # H9268-5G).. This method allowed ∼1300 fold coverage of the mGeCKO library A. After 24 hours, cells were trypsinized, pooled, and seeded in 15 cm plates in media containing puromycin for selection (3 µg/ml for the 4T07 cells and 2 µg/ml for the 4T1 cells) (Wisent # 400-160-EM). Cells were propagated under selection for a week while maintaining a library coverage of at least 700 fold (50 x 10^6^ cells). Then, the first time point (T0) was collected (50 x 10^6^ cells/timepoint to keep the stated fold coverage). The cells were propagated in a similar manner while collecting the consequent timepoints (T1, T2, T3). The collected cells were spin down in 15 ml tubes and kept in -80 °C until they were processed. Every screen was repeated three independent times following the same procedure, where replicates were annotated A, B, and C (as shown in Figure S2). The collected samples were then processed for genomic DNA (gDNA) extraction according to the manufacturer’s protocol (Zymo Research Quick-gDNA MidiPrep Plus kit, #D4075) and prepared for NGS (HiSeq 4000 Illumina) as previously described (18).

### Immunostaining, confocal microscopy, and image analysis

Cells were plated on fibronectin-coated glass coverslips to desired confluence ∼6 hrs with the one exception of CAMA-1 cells that were plated directly on glass coverslips 72 hours prior to staining (VWR # CACB354008). For the 4T07-*Pik3c3* shRNA cells, cells were plated on fibronectin-coated glass coverslips and induced with doxycycline (1 µg/ml) (Sigma # D9891) for 48 hours before proceeding to staining. For amino acid starvation, cells were serum starved (O/N; for 16 hrs), then incubated with EBSS for 2hrs. For re-feeding, 50x MEM amino acids solution was diluted to 1X into EBSS and added to the cells for 30 minutes. Cells were fixed with 3.7% formaldehyde in CSK buffer (100mM NaCl, 300mM Sucrose, 3mM MgCl2, 10mM Pipes pH 6.8) for 15 min. Immunostaining was performed in saponin buffer (TBS supplemented with 1% BSA, 0.01% saponin) using the following antibodies: Rat anti-LAMP1 (Abcam #ab25245) at a concentration 1/400, Mouse anti-LAMP-2 (Developmental Studies Hybridoma Bank #H4B4) at a concentration 1:50, Mouse anti-phospho(Ser2448)-mTOR (Cell Signaling Technology #2971) at a concentration 1:200, and Rabbit anti-Rptor (14C4) (Bioss antibodies, #51285M) at a concentration 1:100. Images were collected from a Zeiss confocal LSM700 (Carl Zeiss, Jena, Germany) microscope with oil immersion objective lenses (Plan-Apochromat, 60×, 1.40 numerical aperture NA; Carl Zeiss).

Image analysis was performed using the FIJI software version 2.3. For measurement of pmTOR at the lysosomes, a region of interest corresponding to the lysosomes was created using the auto-threshold of the LAMP1 channel and was transferred to the corresponding background subtracted pmTOR channel to measure the mean fluorescence intensity (MFI) at the lysosomes. Data represent the mean with SD of at least 30 images from three independent experiments. Statistical significance was determined using one-way ANOVA test with a confidence interval of 95%. The lysosome distribution was measured from confocal images of LAMP1 and pmTOR co-staining captured as described above. For at least 75 cells per condition, a single line, five pixels in width, was traced manually from the center of the nucleus to the cell edge as delineated by pmTOR staining (See Fig. S5). The intensity values along this line were obtained using the “plot profile” plugin from FIJI. The mean fluorescence intensity and the positions along the line for each cell were normalized from 0 to 1 using Matlab as used in reference (19). The normalized data were transferred to Prism 9 software (GraphPad Prism, GraphPad Software, San Diego CA) for further analyses. The percentage of signal at the periphery was calculated by dividing the area under the curve (AUC) between positions 0.75 to 1 by the total AUC between positions 0 to 1. Data are representative of three independent experiments and are shown as the mean with SD (n>75). Statistical significance was determined using the non-parametric Mann-Whitney test with a confidence interval of 95% (****, p-value < 0.0001). This test analysis compares the distributions of two unpaired groups. Data are shown as the mean with SD. For tubulation quantification, the percentage of cells presenting at least one tubulated lysosome was determined among at least 220 cells per condition from three independent experiments. The percentages in the DMSO- and VPS34IN1-treated cells (PDX1915) were then compared by a two-tailed t-test to calculate the *p*-value as stated on the figure.

### Animal Studies

All animal experiments were performed in accordance with the Canadian Council of Animal Care guidelines and approved by the Animal Care Committee of the Montreal Clinical Research Institute (IRCM).

#### Primary tumor and spontaneous metastasis assays

For the 4T07 (Fig. S9) and 4T1 (Fig. S10) experiments,1×10^5^ 4T07-TGL or 4T1-TGL cells were, respectively, injected in the fat pad of the 4^th^ mammary gland of 6 week-old NU/J mice (The Jackson laboratory #002019). For the 4T07 experiment, 1-week post-injection, the mice were randomized into two groups to receive either vehicle (5% DMSO in corn oil) or VPS34IN1 (50 mg/kg/day) by oral gavage daily for either 6 days or 12 days. For the 4T1 experiment (in nude mice), 1-week post-injection, mice were randomized to receive either vehicle or VPS34IN1 (50 mg/kg/day) for two weeks.

For experiments in (Fig. 7A-C), 2×10^5^ 4T07 cells were resuspended in 50 µL of sterile PBS and injected in the fat pad of the 4th mammary gland of 8-10 week-old BALB/C mice (The Jackson Laboratory #0651). On day 24 post-injection, mice received one daily oral gavage of 50 mg/kg/day of VPS34IN1 for 6 consecutive days. On day 30, mice were euthanized to recover the primary tumor and the lungs. Lungs were placed in 2 mL of 10% FBS-supplemented DMEM, chopped into pieces, and incubated with 10 mg/mL collagenase IV (Sigma #C5138) for 3-4 hours at 37°C. The tissue preparation was vortexed every hour to facilitate the tissue digestion. A single cell suspension was prepared by pipetting up and down for 25 times with a P100, and filtered though a 100 µm filter to remove debris. Cells were pelleted to perform a red blood cell lysis according to the manufacturer’s instructions (Sigma #R7757). Half of the lung cell preparation was plated on a 10-cm dish in DMEM supplemented with 10% FBS, 1% P/S, and 25 µM thioguanine (Sigma #A4660), in duplicate. Colonies were allowed to grow for 7 days before staining with Crystal Violet. For the staining with Crystal Violet, plates were washed twice with sterile PBS, fixed for 20 minutes with methanol, stained with a Crystal Violet solution (0.5% Crystal Violet (Sigma #C6158); 25% methanol; 75% PBS) for 40 minutes, and washed twice with sterile water. Plates were allowed to dry overnight and colonies were counted using a light microscope. Results were plotted using GraphPad Prism (GraphPad Software, San Diego CA). Different conditions were compared by Mann-Whitney test to calculate statistical difference and resulting p-value is indicated on the graph.

For experiments in (Fig. 7D-H), 5×10^5^ 4T1 cells were injected in the fat pad of the 4^th^ mammary gland of 8-10 week-old BALB/c mice (The Jackson Laboratory #0651). Two weeks later, primary tumors were resected. Mice were left to recover for ∼3-5 days, then randomized for treatments with either vehicle or VPS34IN1 for a week as described above. At the experimental endpoint, mice with recurred primary tumors were excluded from the final analyses presented in Figure 7. Otherwise, mice were sacrificed and lungs were collected immediately, emersed in PBS then fixed in 4% PFA for 24 hr at 4°C. Lungs were then placed in 70% ethanol before being embedded in paraffin. Paraffin blocks were then sectioned at 4 μm thickness and processed for H&E stainings and the 20X objective of the Aperio-XT slide scanner (Leica Biosystems) was used to scan slides. Fiji (ImageJ) was used to visualize the generated images and measure size of metastatic lesions. Given the irregularity between different lesions, the length of the longest dimension was considered as a measure to categorize micrometastases (< 100 um), macrometastases (> 200 um) and lesions scoring in between (100-200 um). The vehicle and VPS34IN1-treated groups were compared by a two-tailed t-test to calculate the *p*-value (n= 6 or 7 mice/group).

#### Lung Disseminated tumor cells (DTCs) burden in 4T07 tumors-bearing nude mice (early timepoint)

At the experimental endpoint, mice were euthanized, and lungs were processed as previously described (6). Briefly, lungs were dissected and immediately emersed in PBS and cut into small pieces. Then, the lung pieces were digested in PBS supplemented with 75 µg/ml TH Liberase (Roche #05401127001), 75 µg/ml TM Liberase (Roche #05401151001), and 12.5 µg/ml DNAse. Samples were then incubated in the digestion buffer on a rotor in 37 °C. Samples were then centrifuged at 1,300 RPM for 5 minutes and supernatant was discarded. Pellets were resuspended in PBS, pipetted up and down multiple times and filtered through a 70 µm strainer to get rid of the undigested lung pieces. 1 ml of Red Blood Cells Lysis Solution (Sigma #R7757-100ML) was then added to each sample and the manufacturer protocol was followed. Cells were finally seeded in 15 cm plates in Thioguanine-containing DMEM Wisent media supplied with 10% FBS and 1 % P/S antibiotics. Plates were left 48 hours and d-luciferin (Promega #E1605; dissolved in PBS) was added to the DMEM media in a concentration (150 µg/mL). Plates were then imaged using the Xenogen IVIS 200 machine (PerkinElmer) and analyzed with Living Image 4.2 software.

The difference in the DTCs burden between the vehicle- and VPS34IN1-treated groups was quantified as a foldchange difference in bioluminescence emitted signal between the two groups. Foldchange values were represented in Log values and the two groups (control and drug treated) were compared by a two-tailed t-test to calculate the *p*-value. The resulting values are stated on the graph and in main text.

#### Ex vivo metastasis assay

Mice were injected intraperitoneally with 150 mg/kg d-luciferin (in PBS) solution and were euthanized 5-7 minutes later. Lungs were immediately collected and imaged using the Xenogen IVIS 200 machine (PerkinElmer).

### Two-color competition assays

Individual gRNAs targeting genes, or the Rosa locus (sequences provided below) were cloned in either Lentiguide-gRNA-NLS-GFP-2A-PURO or Lentiguide-NLS-mCherry-2A-PURO using the golden gate assembly method (18,20). Producing lentiviruses from the cloned vectors and the LentiCas9-Blast (Addgene #52962) was performed as detailed earlier but with scaling down the production procedure. The 4T1 or the 4T07 cells were first transduced with the LentiCas9-Blast and selected with blasticidin (5 µg/ml) (ThermoFisher Scientific # R210-01). An MOI test for the produced individual gRNAs lentiviruses was performed. The 4T1-Cas9 and 4T07-Cas9 cells were subsequently infected and selected with puromycin as detailed earlier.

Immediately after selection, 5 x 10^4^ of the 4T1-gRNA (in Lentiguide-gRNA-NLS-GFP-2A-PURO) were seeded with 5 x 10^4^ of the 4T1-Rosa gRNA (in Lentiguide-NLS-mCherry-2A-PURO) in 12 well-plate and left overnight. The following day, the plates were imaged using the 4x objective of IncuCyte (Essen Bioscience) to assess the representation of the green and red populations at the 1^st^ timepoint of the assay (Passage 0, P0) by counting the GFP- and mCherry-positive nuclei. Every two days, cells were split 1:10 in new 12 well plates, kept overnight and imaged the following day to assess the two populations representation. This pipeline was repeated over 10 passages. The same procedure was repeated for all the investigated gRNAs (targeting *Pik3ca*, *Pik3c3*, and *Pik3r4*) in this study.

In every passage, the percentage of the GFP-positive population from the total population (GFP- and mCherry-positive combined) was calculated. Then the ratio between the percentage of the GFP-positive population at every passage over passage 0 (reference timepoint) was used to assess the dropping out of the GFP-positive population. The calculated ratio in the last passage (P10) of each condition (i.e. gRNA) was compared to that of the control condition (i.e. the two red and green co-cultred populations are expressing gRNA targeting the ROSA locus) by a two-tailed t-test to calculate the *p*-value. The resulting values are stated on graphs.

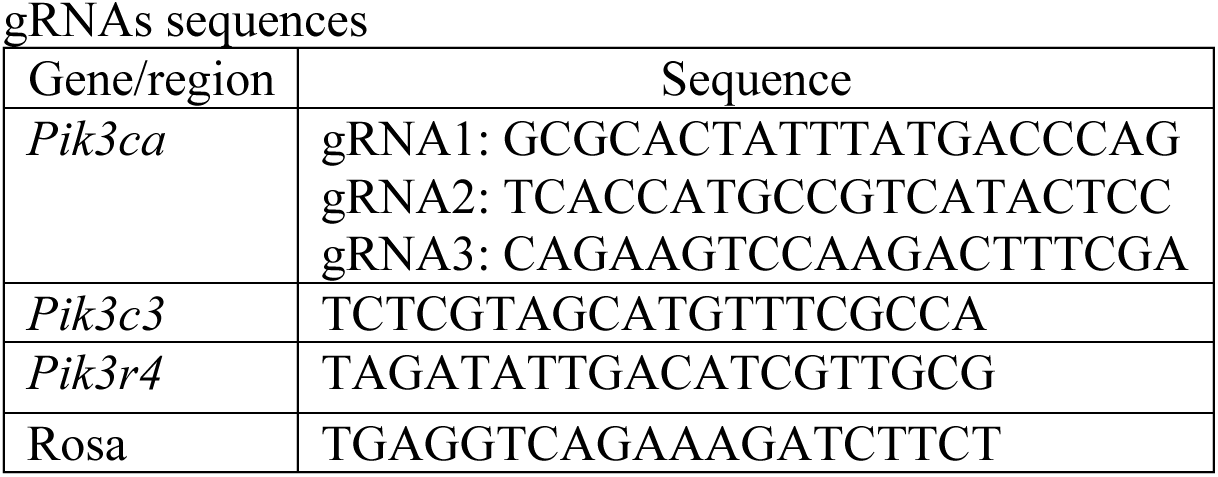

### Proliferation assays

5 x 10^3^ 4T1 or 4T07 cells were seeded in 48-well plate O/N and the corresponding drugs (BYL719 or MK-2206) were added in different concentrations (2 wells/concentration as technical replicates). Plates were immediately transferred to be imaged with the IncuCyte (Essen Bioscience). After 72 hours, the cells’ confluency was determined for all the drug concentrations. The increase in confluency was determined for every condition in reference to the confluency at the first timepoint (immediately after adding the drugs). Then a ratio between the determined change in confluency under different drug doses to that of the control condition (DMSO-treated) was calculated for each cell line. These values for the two cell lines were then plotted to compare the response of the two cell lines to the same drug. The values (response to same dose in reference to DMSO treated condition) of the two cell lines were compared by a two-tailed t-test to calculate the *p*-value. The resulting values are stated on graphs.

### 3D culture and cell death assessment

This 3D in vitro culture conditions have been used as a surrogate for in vivo extracellular matrix environment. To mimic matrix-associated growth factors, 96-well cell culture plates were coated with 40 mL Cultrex® Basement Membrane Extract (BME) (21) (R&D #3433-010-01), containing laminin, collagen IV, entactin, and heparan sulphate proteoglycan and left to polymerize to hydrogel for 30 minutes at 37 °C. Three thousand of 4T1 and 4T07 cells suspended in RPMI-1640 media supplemented with 2% FBS, 2% BME and 1% P/S were seeded onto BME. Plates were further placed into IncuCyte imaging system (Essen Bioscience) and imaged once every 12 hours with the 10x objective. Four days post seeding, the VPS34IN1 drug was added at the indicated doses (5 uM or 10 uM) together with SYTOX™ Near-IR Dead Cell Stain (ThermoFisher Scientific #S11382) and imaged for the following 4 days to quantify the cell death burden. At the experimental endpoint (8th day post seeding), counts positive for SYTOX™ Near-IR were quantified via the IncuCyte software and data was plotted in GraphPad Prism (GraphPad Software, San Diego CA). Different conditions were compared by two-way ANOVA test and p-values are indicated on the graphs.

### Protein extraction, western blotting, and activity analysis

Cells were rinsed with PBS and lysed using ice-cold Radio ImmunoPrecipitation Assay (RIPA) buffer (1% Nonidet P-40, 150 mM NaCl, 50 mM Tris pH 7.5, 5 mM EDTA, 0.5% deoxycholic acid, 0.1% SDS) supplied with 1mM Na3VO4, 5 mM NaF and 1× complete protease inhibitor (Roche, Indianapolis, IN). Samples were then centrifuged (15 min, 13,000g, 4°C) and boiled with 6x Laemmli sample buffer for 5 min at 95 °C for denaturation.

Tumors excised from mice were placed in a mortar filled with liquid nitrogen and then mechanically disrupted with a pestle. RIPA buffer (prepared as stated earlier) was added to the disrupted tumor pieces. Once the buffer returned to the liquid state, the mix of the buffer with the tumor pieces were transferred to tubes and pipetted vigorously up and down several times. Tubes were then left on ice for 15 minutes. Samples were centrifuged, and supernatants were transferred to new tubes and boiled with 6x Laemmli sample buffer as detailed earlier.

Protein separation was then performed using the standard SDS-PAGE protocol and proteins were transferred to nitrocellulose membranes (BioRad #1620115) via wet transfer for 2.5 hours at 4°C. Membranes were subjected to blocking for 0.5-1 hour in Tris-Buffered Saline (TBS) solution supplemented with 0.1% Tween-20 (Bio Basic #TB0560) (TBS-T) and 1% bovine serum albumin (Wisent #800-095-CG) before incubation with primary antibodies O/N at 4°C. The following day, membranes were washed (3 x 10 minutes) in TBS-T and incubated with secondary antibodies for 1 hour at room temperature. Membranes were washed (3 x 5 minutes) in TBS-T and incubated with Enhanced Chemiluminescence substrate (ECL; BioRad #1705061) for 1 minute prior to exposure.

All primary antibodies were diluted in TBS-T supplemented with 1% BSA and used as per the following concentrations: rabbit anti-phospho-AKT(S473) (#9271S), rabbit anti-AKT (#9272S), rabbit anti-phospho-S6K1 (Thr389) (#9205S), rabbit anti-S6K1 (#9202S), rabbit anti-phospho-S6 (Ser240/244) (#2215S), rabbit anti-S6 (#2217S) were obtained from Cell Signaling Technology and used at a concentration (1:1000). Mouse anti-α-Tubulin (Sigma Aldrich #T5168) was used at a concentration (1:8000). Secondary antibodies: Mouse anti-rabbit IgG-HRP (Santa Cruz #2357) for pAKT, AKT, pS6K, S6K, pS6, and S6 was diluted (1:5000) in TBS-T supplemented with 1% BSA. Goat anti-mouse IgG-HRP for α-Tubulin (Sigma Aldrich #A4416) was used at the same concentration. An anti-FLAG M2-peroxidase (HRP; Sigma #A8592) was used at a concentration of 1:8000 in TBS-T supplemented with 1% BSA to assess the expression of PIK3C3 as shown in Figure S7.

The gel analyzer tool of Fiji (ImageJ) was used to quantify the western blot signals. mTORC1 activity was assessed as a ratio between the pS6K and total S6K signals. The calculated ratio (i.e. mTORC1 activity in arbitrary units) was either used directly to compare the 4T1 and 4T07 cells (ex: Fig 3A-C) or normalized to show relativety between the two lines (ex: Fig S5A-C). In all cases, a two-tailed t-test was used to calculate the *p*-value for the difference between the tested two conditions. The calculated *p*-values were stated in figures and text. To quantify the mTORC1 activity in the analyzed tumors, the phosphoS6K1/total S6K1 ratio was similarly calculated for every sample as a readout. Then, to compare two groups (drug vs vehicle) considering the variability in activity even in the same group, the foldchange of each sample’s activity was calculated relatively to each sample’s activity in the opposing group and an average was determined as a final value for each sample. The latter calculation was made for every sample in the two groups. Finally, the resulting values were represented in log values and the two groups (control and drug treated) were compared by a two-tailed t-test to calculate the *p*-value. The resulting values are stated on the graphs.

## RESULTS

### Genome-wide CRISPR screens define differential fitness genes in 4T1 and 4T07 cells

We characterized the molecular subtype of BC modeled by 4T1 and 4T07 cells. We generated and correlated RNA-Seq transcriptomic data from these two cell lines with gene expression data from 47 human BC cell lines. We conclude that 4T1 and 4T07 cells are closer to the human basal B molecular BC subtype predominantly within the triple negative clinical BC subtype, complementing a recent investigation of 4T1 cells (22) (Fig. S1; Table S1). The 4T07 cells display features of the basal B subtype consistent with a common origin of the 4T1 and 4T07 cells (Fig. S1; Table S1).

Genome-wide loss-of-function CRISPR screens identify core and context-dependent fitness genes, revealing common and specific active signaling nodes (13,23,24). Fitness genes are those that when perturbed result in a proliferative or survival disadvantage for cells. We hypothesized that fitness genes specific to 4T1 may be involved in survival or metastatic progression, while those specific to 4T07 might regulate survival during dormancy or outgrowth when implanted in permissive microenvironments. To explore this, we used a CRISPR screening pipeline on 4T1 and 4T07 cells, employing the mGeCKO library encompassing ∼70,000 gRNAs targeting ∼21,000 murine genes (18). Following selection for cells with gRNA integration (T0), genomic DNA was extracted at 3 consecutive timepoints over 3 weeks (T1, T2, and T3) to map gRNAs representation relative to T0 (Fig. 1A). Quality control analyses confirmed adequate library coverage (Fig. S2A-B) and clustering of biological replicates (Fig. S2C-D). The two screens showed high sensitivity, with precision-recall effectively distinguishing essential from non-essential genes (Fig. S2E-F). Finally, fold-change distributions of gRNAs targeting essential genes, over the course of the screens, were prominently shifted while those targeting nonessential genes showed minimal change (Fig. S2G-H). These analyses demonstrated the screens’ robustness at identifying fitness genes in both cell lines.

**Figure 1.**
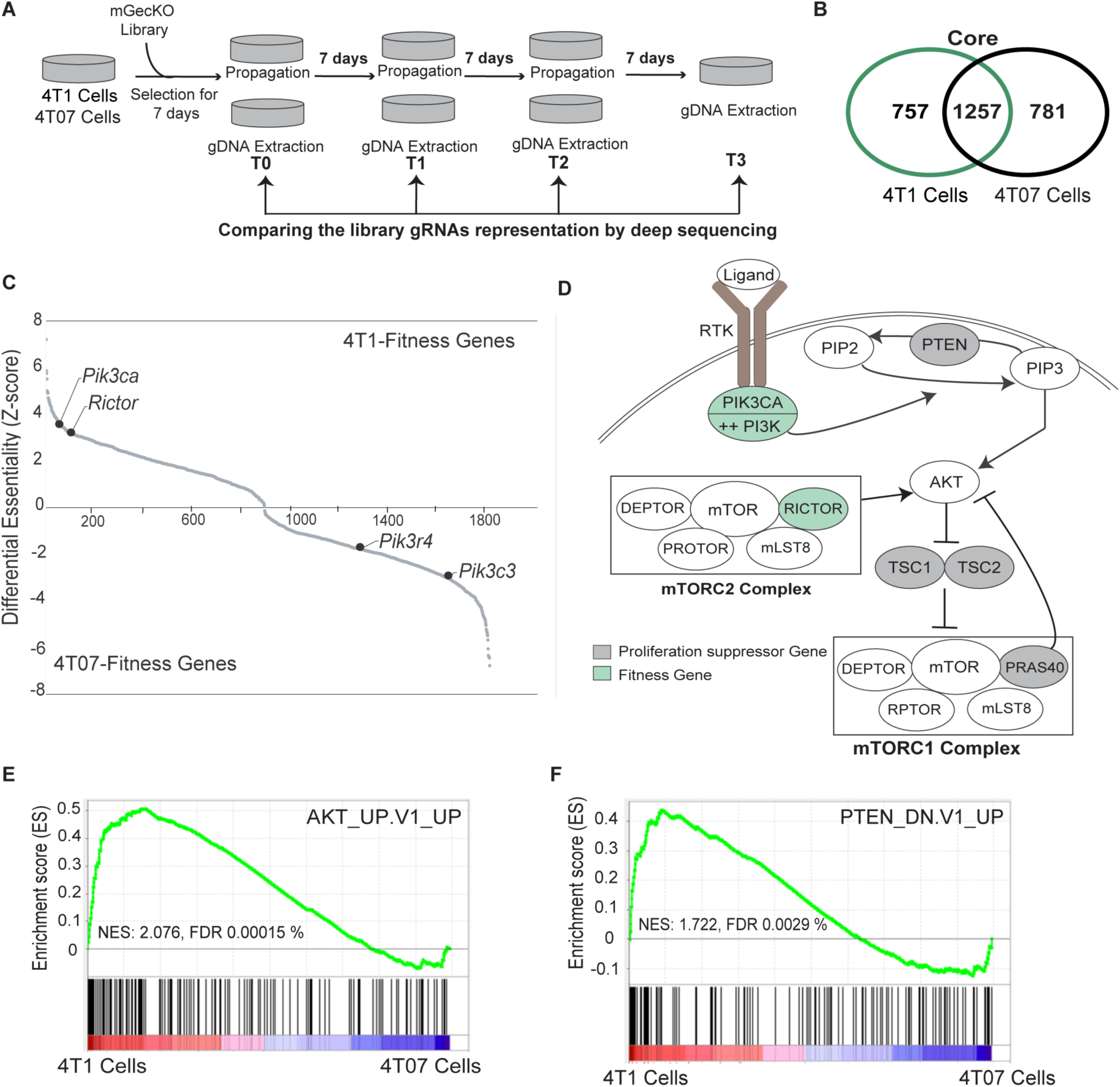
Genome-wide knockout CRISPR screens define the common and differential fitness genes in the 4T1 and 4T07 cells. **(A)** Schematic outline for the performed screens’ pipeline. **(B)** Enumeration of the revealed cell line-specific and common (core) fitness genes. **(C)** Ranked differential gene fitness score between the 4T1 and 4T07 cells. Value > 0 denotes 4T1-specific fitness gene; value < 0 denotes 4T07-specific fitness gene. **(D)** Schematic representation integrating the 4T1-specific fitness (in green) and proliferation suppressor genes (positively selected genes; in grey) in the canonical PI3K-AKT pathway. **(E-F)** GSEA analysis comparing the enrichment of genes previously found to be induced by AKT upregulation **(E)** or PTEN downregulation in the two cell lines **(F)**.

We identified 2014 fitness genes in 4T1 cells and 2038 in 4T07 cells, including 1257 as common genes (Table S2) that functionally mediate biological processes known for core essential genes in cancer (Fig. 1B; Fig. S3) (13,25). Consequently, 757 and 781 specific fitness genes were identified in 4T1 and 4T07 cells, respectively (Fig. 1B; Table S3-4). We focused on identifying differential signaling pathways dependencies. Among the top ranking 4T1-specific fitness genes were *Pik3ca* (catalytic subunit of class-I PI3K) and *Rictor* (core component of mTOR complex 2 (mTORC2)) (Fig. 1C), both being positive regulators of the PI3K pathway (Fig. 1D). Conversly, PI3K negative regulators like *Pten*, *Tsc1*, *Tsc2*, and *Pras40* were identified as proliferation suppressor genes, whose gRNAs were enriched at the last timepoint (T3) in 4T1 cells (Fig. 1D and Table S5). These data suggested that 4T1 cells may possess higher PI3K-AKT activity in comparison to 4T07 cells. Indeed, a GSEA analysis of the two cell lines’ global transcriptome identifed elevation of *AKT*– and downregulation of *PTEN* gene signatures to be enriched in 4T1 in comparison to 4T07 cells (Fig. 1E-F). By analyzing the 4T07-specific fitness genes, we identified *Pik3c3* (catalytic subunit of class-III PI3K) and its regulatory subunit *Pik3r4* (Fig. 1C). Given the clinical relevance of class I PI3K pathway and the less explored class III PI3K pathway in BC, we investigated these differential dependencies.

### Validation of *Pik3ca* and *Pik3c3* as differential fitness genes for 4T1 and 4T07 cells

To confirm the essentiality of *Pik3ca* in 4T1 cells, we performed a two-color competition assay (20). We generated 4T1 cells expressing Cas9 and then infected them with either a vector to express a *Pik3ca*-targeting gRNA and GFP or a vector to express mCherry and a gRNA targeting *Rosa26* (Fig. 2A). Co-culturing these lines in a 1:1 ratio led to the dropout of cells expressing *Pik3ca*-targeting gRNA(s) as indicated by monitoring the percentage of green cells within the total population over 10 passages (Fig. 2B-C). We validated these findings pharmacologically by treating 4T1 and 4T07 cells with the PIK3CA selective inhibitor BYL719 (26), which significantly inhibited 4T1 cell proliferation but did not affect 4T07 cells (Fig. 2D). Next, we investigated the effect of the AKT inhibitor MK-2206, and found that it inhibited the proliferation of 4T1 cells more prominently than 4T07 cells (Fig. 2E).

**Figure 2.**
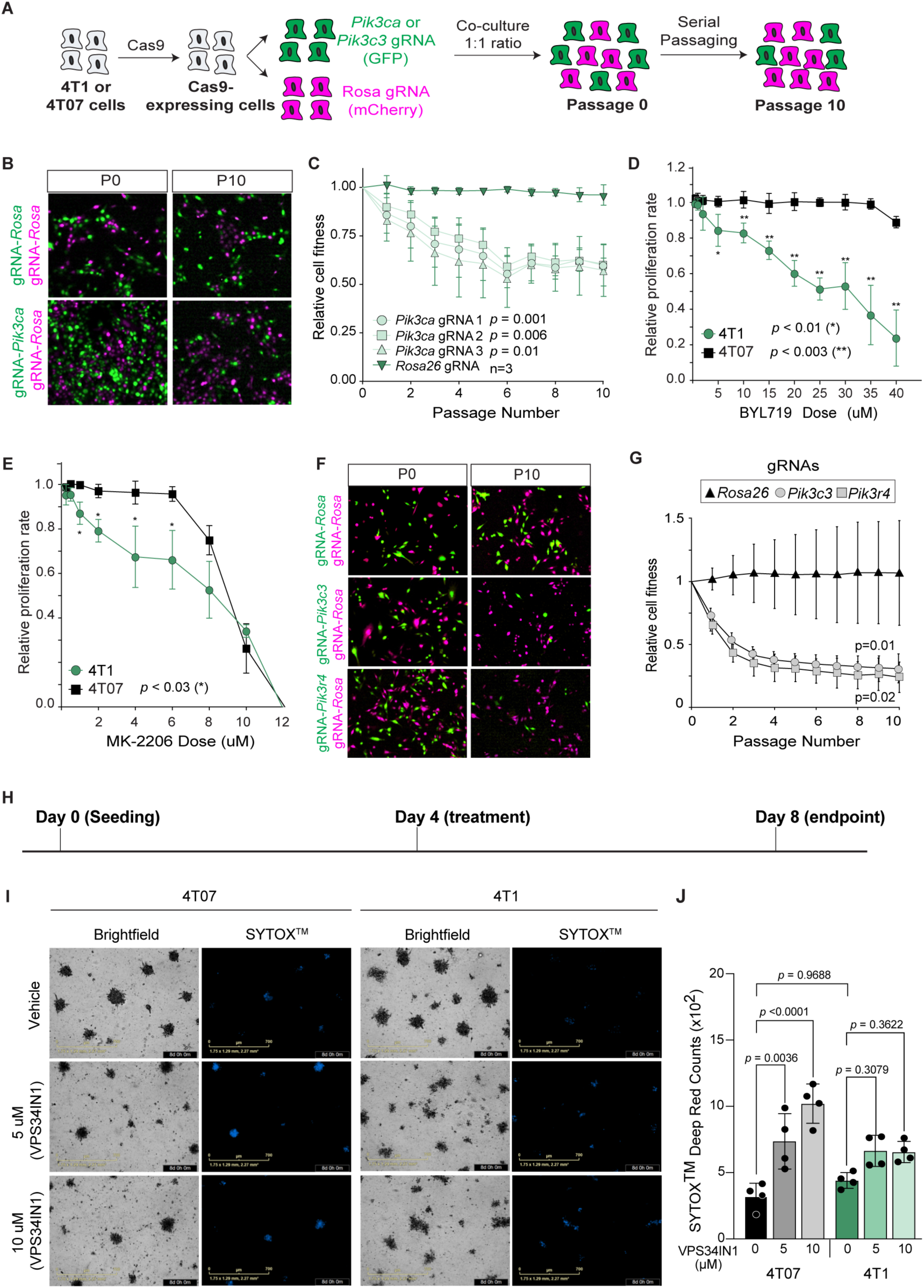
*Pik3ca* and *Pik3c3* are differential fitness genes for the 4T1 and 4T07 cells. **(A)** Schematic representation for the two-color competition assay. **(B)** Representative images showing either 4T1 cells expressing *Pik3ca*-gRNA or *Rosa26*-gRNA (both in green) co-cultured independently with 4T1 cells expressing *Rosa26*-gRNA in magenta, at passages zero (P0) and ten (P10). **(C)** Graph showing representation of the green 4T1 cell populations over serial passages in reference to P0. P values indicate difference between control condition and experimental conditions at P10. **(D)** Proliferation assay comparing the 4T1-4T07 cells’ response to BYL719. **(E)** Proliferation assay comparing the 4T1-4T07 cells’ response to MK-2206. **(F)** Representative images showing either 4T07 cells expressing *Pik3c3*-gRNA, *Pik3r4*-gRNA, or *Rosa26*-gRNA (all in green) co-cultured independently with cells expressing *Rosa26*-gRNA in magenta at passages zero (P0) and ten (P10). **(G)** Graph showing representation of the green 4T07 cell populations over serial passages in reference to P0. P values indicate difference between control condition and experimental conditions at P10. **(H)** Schematic for the timeline of assessing the effect of VPS34IN1 on the 4T07 and 4T1 cells’ survival in (I-J). 4T07 and 4T1 cells were seeded and cultured on basal membrane extract (BME) where they form spheroids, and were then treated with VPS34IN1 or DMSO. **(I)** Representative images of 4T07 and 4T1 3D spheroids at the experimental endpoint after treatment with either the vehicle (DMSO) or VPS34IN1 at 5 uM and 10 uM. **(J)** Quantifications of cell death burden as indicated by positive staining for SYTOX^TM^.

To validate the essentiality of *Pik3c3* in 4T07 cells, we employed the same two-color competition assay (Fig. 2A). We observed a significant drop in the population of 4T07 cells expressing gRNAs targeting *Pik3c3* or *Pik3r4* over time, as compared to the control (Fig. 2F-G). In contrast, in a similar assay, 4T1 cells showed only modest reduction (Fig. S4A-B). Comparing the representation of *Pik3c3* gRNA-expressing 4T1 and 4T07 populations after normalization to their respective control conditions confirmed the predicted differential dependence on *Pik3c3* (Fig. S4C). To validate these findings pharmacologically, we treated 4T07 and 4T1 spheroids with two doses of VPS34IN1, a PIK3C3-specific inhibitor (27), and assessed cell death (Fig. 2H). VPS34IN1 treatments led to a 2-to 3-fold increase in death rate of 4T07 cells, with no significant effect on the 4T1 cells, compared to vehicle-treated controls (Fig. 2I-J). These findings indicate that *Pik3c3* is essential for 4T07, but not 4T1, cell survival .

These data demonstrate that the diffential genetic dependence on PI3K classes in 4T1 and 4T07 cells is reflected in differential vulnerability to specific inhibitors of these PI3Ks. This prompted us to investigate whether these phenotypes result from differential signaling activity in the PI3K-AKT-mTORC1 pathway.

### mTORC1 is more active in 4T07 cells compared to 4T1 cells

We assessed the activity of downstream components of the PI3K pathway. 4T1 cells exhibited higher PI3K activity in untreated conditions, indicated by increased phosphorylated Akt levels (Fig. 3A-C). Short-term treatments with BYL719 similarly inhibited the Pi3k-Akt-mTORC1 pathway in both cell lines, as demonstrated by reduced levels of phosphorylated Akt, S6k1, and S6 (Fig. 3A). However, short-term Akt inhibition did not mediate the same effects as Pik3ca inhibition (Fig. 3B). Specifically, mTORC1 activity, measured by phospho-S6k1 levels, was reduced only in 4T1 cells upon Akt inhibition. Prominently in the untreated conditions in both experiments (Fig. 3A-B), 4T07 cells demonstrated a two-fold higher mTORC1 activity than 4T1 cells (Fig. 3C), despite the dispensibility and low activity of the class I Pi3k in 4T07 cells (Fig. 3A-C). This finding is intriguing from a signaling and biological perspective. First, sustained or reactivated mTORC1 activity confer resistance to PIK3CA inhibition (28,29). Second, PIK3C3 mediates mTORC1 activity especially when class I PI3K is inhibited long-term (30). Third, high mTORC1 activity is known to support survival of dormant cells *in vivo* (31). This led us to further explore the regulation of mTORC1 activity in 4T1 and 4T07 cells.

**Figure 3.**
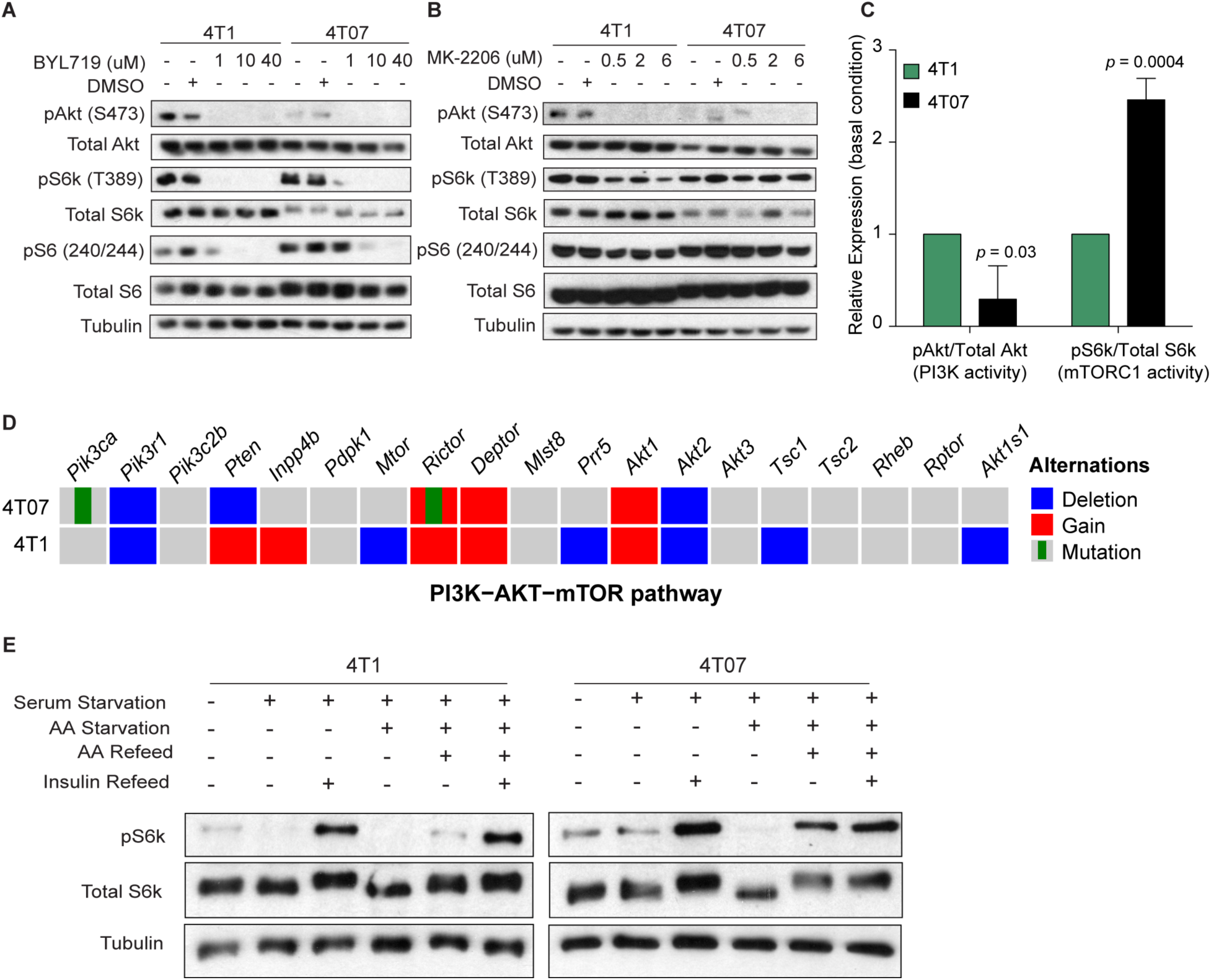
mTORC1 is more active in 4T07 cells compared to 4T1 cells. **(A-B)** Western blot analysis for the Pi3k-Akt-mTORC1 pathway readouts in the 4T1-4T07 cells upon treating with BYL719 **(A)** or MK-2206 **(B)**. **(C)** Quantification of the relative PI3K and mTORC1 activity in the two cell lines from the performed western blots in (A) and (B). **(D)** Oncoprints for somatic alterations in genes involved in the PI3K-AKT-mTOR pathway in 4T1 and 4T07 cells. **(E)** Western blot analysis assessing mTORC1 activity in the 4T1-4T07 cells under different nutritional conditions. AA; amino acids.

We investigated the relience of each cell line on Pi3kca for mTORC1 activity. Low-dose BYL719 (1 uM), under serum starvation or subsequent insulin stimulation, reduced mTORC1 activity preferentially in 4T1 over 4T07 cells (Fig. S5A-C), indicating that 4T07 cells are less dependant on Pi3kca for mTORC1 regulation. To determine whether mTORC1 hyperactivity in 4T07 cells affects their response to BYL719, we treated 4T07 cells with a high dose of BYL719 (10 uM; did not impact proliferation (Fig. 2D)) and monitored mTORC1 activity over time. We observed that mTORC1 activity started to increase after 48 hours of treatment (Fig. S5D). These findings suggest non-canonical regulation mTORC1 in 4T07 cells.

To determine whether genomic alterations in PI3K-AKT-mTORC1 pathway genes could explain the differences between 4T1 and 4T07 cells, we performed whole-genome sequencing (WGS) on both cell lines, in reference to genomic DNA from a female BALB/c mouse. Despite high sequencing coverage (see Supplementary Methods), we found no significant differential genomic alterations of established functional consensequences in the Pi3k-Akt-mTORC1 pathway genes (32) to explain the differential dependence and signaling activity (Fig 3D). We identified two missense mutations in 4T07 cells: *Pik3ca* (p.A423S) and *Rictor* (p.R576S) (Fig. 3D; Table S6). Both mutations occur in conserved residues and are predicted to have moderate impact, but they have not been found in human tumors (n= 75,661 patients) (see Supplementary Methods), suggesting limited functional relevance. We also noted differential copy number alterations in *Pten*, *Inpp4b*, *Mtor*, *Prr5*, *Tsc1*, and *Akt1s1* (Fig. 3D). Only alterations in *Inpp4b, Tsc1,* and *Akt1s1* correlated with mRNA expression patterns between the two lines (Tables S7-S8). Interestingly, 4T1 cells demonstrated genomic gain, correlated with mRNA overexpression, of *Inpp4b,* a tumor suppressor and a negative regulator of PI3K activity that is frequently lost in basal BC (33,34) (Fig 3D; Table S8). Conversely, 4T1 cells showed losses in *Tsc1* and *Akt1s1,* which align with their indentification as proliferation suppressors of these cells in our CRISPR screens (Figure 1D). Although these results suggest that 4T1 cells are wired to restrict the negative regulators of the canonical Pi3k pathway, none of these could explain how mTORC1 activity is elevated in 4T07 cells.

mTORC1 activity is regulated by both growth factors and amino acids (29). Growth factors act mainly through PIK3CA-AKT (29), while amino acids act through various mediators, including PIK3C3 (35). Hence, we examined mTORC1 activity under different nutritional conditions to identify key regulatory mechanisms in 4T07 cells. Compared to 4T1 cells, 4T07 cells exhibited higher basal mTORC1 under serum-fed conditions and maintained residual mTORC1 activity under starvation (Fig. 3E). Amino acid starvation following overnight serum starvation completely abbolished mTORC1 activity in 4T07 cells (Fig. 3E), while amino acid refeeding in serum-free conditions restored the activity to baseline levels (Fig. 3E). Similarly, amino acid refeeding induced the mTORC1 activity in 4T1 cells. These observations suggest differential regulation of mTORC1 in the two cell lines and highlight the potential significance of the amino acid signaling in 4T07 cells.

### Differential mTORC1 activity is reflected in differential lysosomal positioning in 4T1 compared to 4T07 cells

In addition to stimulating mTOR tethering on lysosomal surfaces to allow its consequent activation (29), amino acids activate PIK3C3. The latter mediates mTORC1 activity through different mechanisms, like lysosomal positioning, where mTORC1 activity is higher when lysosomes are positioned in in the periphery compared to perinuclear (35–38). This led us to investigate lysosomal mTORC1 levels and lysosomal positioning in 4T1 and 4T07 cells.

We assessed the co-localization of Rptor (core component of mTORC1) and the phospho-S2448 residue of the mTOR (more specific to mTORC1 than mTORC2 (39)) with the lysosomal marker Lamp1 (Fig 4A-B). Under basal conditions, we observed a diffuse staining pattern for pMtor, with colocalization with Lamp1-positive lysosomes in both cell lines (Fig 4A-B, 1^st^ row). Amino acid starvation localized mTORC1 to small, diffuse cytoplasmic puncta (40) (Fig. 4A-B, 2^nd^ row), and decreased lysosomal mTORC1 levels in both cell lines (Fig. 4A-C). These effects were rescued to baseline by amino acid refeeding (Fig. 4A-C, 3^rd^ row). We also found Rptor in the cytoplasm, where it co-localizes with Lamp1, and the nucleus (Fig. 4A-B). However, the intact mTORC1 was primarily localized in the cytoplasm in both 4T1 and 4T07 cells, with no pMtor signal in the nucleus, consistent with findings in other cells (41,42). Notably, 4T07 cells exhibited 1.4-fold higher levels of lysosomal mTORC1 than 4T1 cells under fed conditions (Fig. 4C). We also observed distinct lysosomal positioning: in 4T07 cells, Lamp1-positive lysosomes accumulate at the tip of cell extensions, while in 4T1 cells, they accumulated in the perinuclear region (Fig. 4A-B, 4D; Fig. S5E-G). This peripheral positioning remained unchange in 4T07 cells by manipulating the nutritional conditions that affected the mTORC1 lysosomal co-localization (Fig. 4D). These results suggest that lysosomes in 4T07 cells are preferentially positioned at the cellular periphery compared to 4T1 cells, providing mechanistic insights into their differential mTORC1 activity.

**Figure 4.**
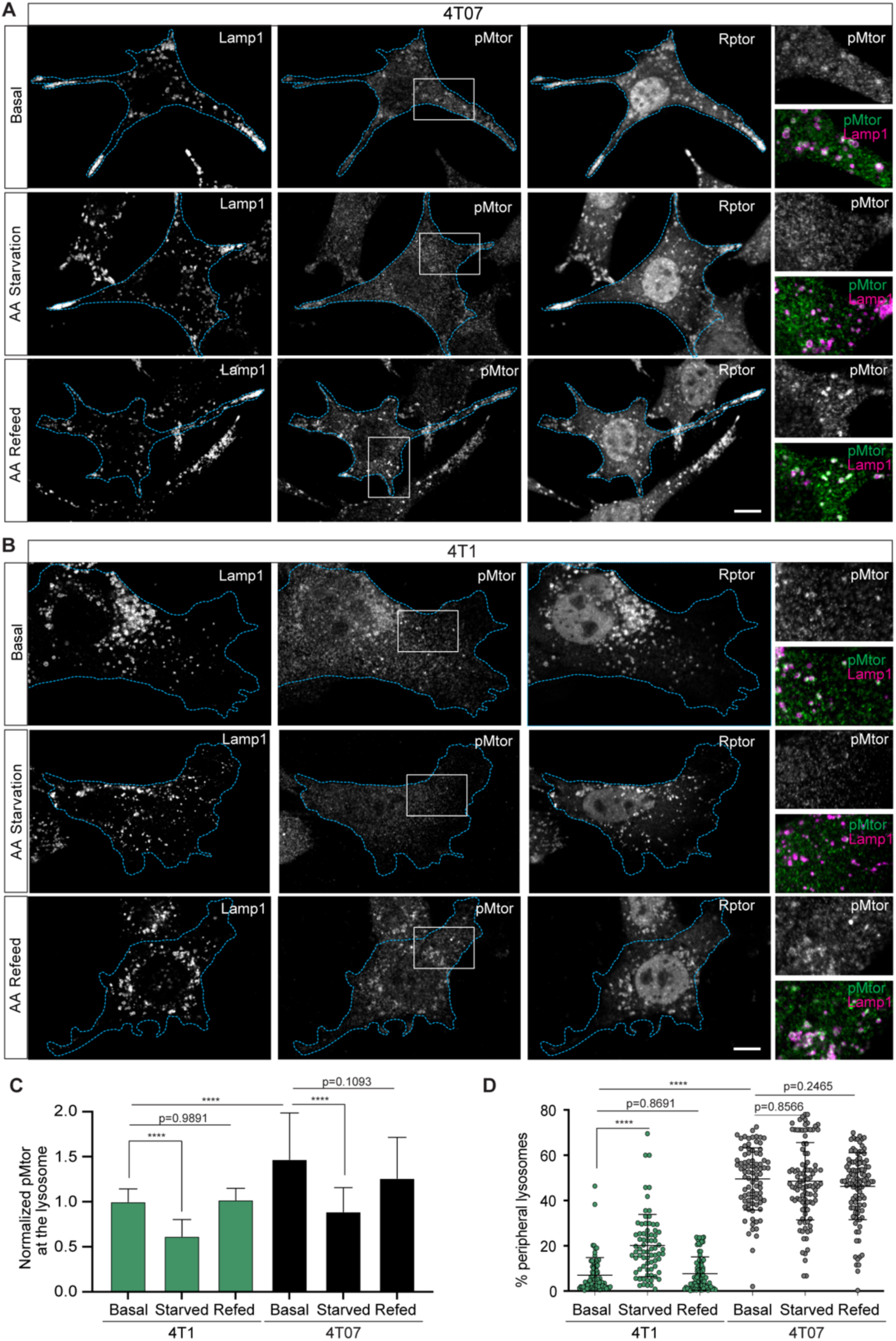
Differential lysosomal positioning between the 4T1 and 4T07 cells. **(A-B)** Immunostaining for the lysosomal marker, Lamp1, and mTORC1 markers, Rptor and pMtor (S2448) in the 4T1-4T07 cells under basal culture conditions or amino acid starvation or amino acid refeeding. **(C)** Quantifications of the mean fluorescence intensity (MFI) of mTORC1 (pMtor signal) on the lysosomes in the two cell lines under the 3 different conditions. **(D)** Graph representing the peripheral lysosomes’ percentage in the two cell lines under the 3 different conditions.

### PIK3C3 mediates peripheral lysosomal positioning and mTORC1 activity in murine and human breast cancer

We investigated whether Pik3c3 regulates mTORC1 activity and/or lysosomal positioning in 4T1 and 4T07 cells. Treatement of 4T07 cells with 1 uM VPS34IN1 for two hours reduced mTORC1 activity by 50% (Fig. 5A-B). A similar trend was observed in 4T1 cells, though with higher variability due to low basal level of mTORC1 activity (Fig. 5A-B). Notably, VPS34IN1 blocked the response of 4T07 cells to amino acid refeeding after starvation (Fig. S6A), aligning with known functions of PIK3C3 (35–37). We then examined the impact of VPS34IN1 on lysosomal positioning and noticed a 10-fold shift of lysosomes from the periphery to the perinuclear area, without affecting mTORC1 tethering to the lysosomes in 4T07 cells (Fig. 5C-D). In 4T1 cells, VPS34IN1 further reduced the basal percentage of peripheral lysosomes by ∼3 fold compared to the control (Fig. 5C-D). To confirm that these effects were specific to Pik3c3 and not due to off-target actions of VPS34IN1, we generated 4T07 cell lines with doxycycline-inducible Pik3c3-targeting or scrambled shRNAs (Fig. S6B). Inducing the expression of these shRNAs demonstrated that the knockdown of *Pik3c3* also shifts the lysosomes from the periphery to the perinuclear region (Fig. S6C-D). These results demonstrate that Pik3c3 regulates mTORC1 activity and lysosomal positioning in both 4T1 and 4T07 cells, with varying magnitude due to differences in basal mTORC1 activity.

**Figure 5.**
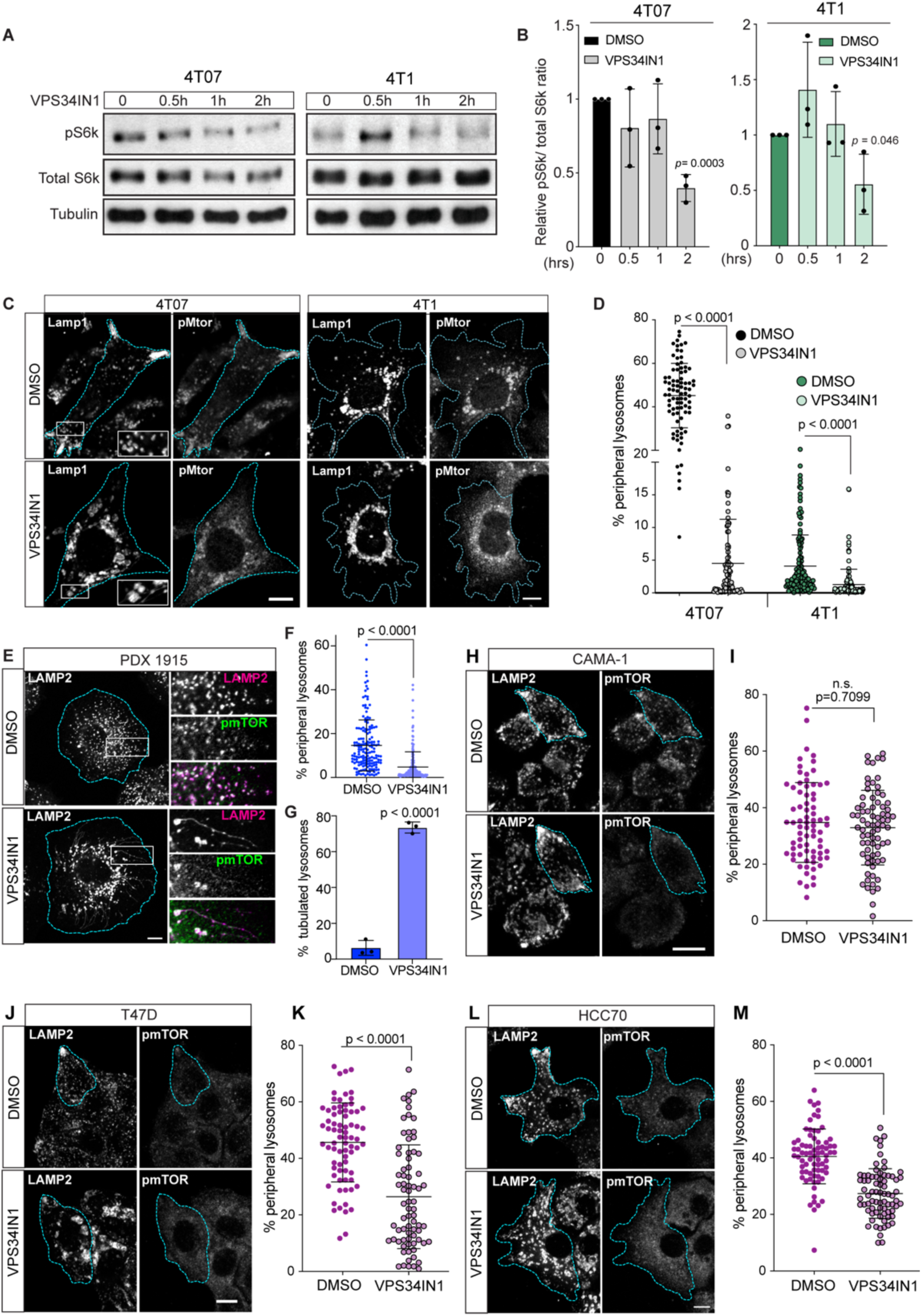
PIK3C3 mediates peripheral lysosomal positioning and mTORC1 activity in murine and human breast cancer cells. **(A)** Western blot analysis for the VPS34IN1 (1 µM) effect on 4T07 and 4T1 cells’ mTORC1 activity. **(B)** Quantification for mTORC1 activity in panel (A). **(C)** Immunostaining for Lamp1 and pMtor in 4T07 and 4T1 cells under the effect of VPS34IN1 (1 µM for 2 hours). **(D)** Quantifications of the percentage of peripheral lysosomes in the VPS34IN1-treated cells in comparison to the control condition (DMSO-treated). **(E)** Immunostaining for LAMP2 and pmTOR in the PDX-1915 cells under the effect of VPS34IN1 (2 uM) or DMSO for 2 hours. **(F)** Quantifications of the percentage of peripheral lysosomes in (E). **(G)** Quantifications of the percentage of tubulated lysosomes in (E). **(H)** Immunostaining for LAMP2 and pmTOR in the CAMA-1 cells under the effect of VPS34IN1 (2 µM) or DMSO for 2 hours. **(I)** Quantifications of the percentage of peripheral lysosomes in (H). **(J)** Immunostaining for LAMP2 and pmTOR in the T47D cells under the effect of VPS34IN1 (2 µM) or DMSO for 2 hours. **(K)** Quantifications of the percentage of peripheral lysosomes in (J). **(L)** Immunostaining for LAMP2 and pmTOR in the HCC70 cells under the effect of VPS34IN1 (2 µM) or DMSO for 2 hours. **(M)** Quantifications of the percentage of peripheral lysosomes in (L).

Next, we assessed whether VPS34IN1-mediated suppression of mTORC1 activity could sensitize 4T07 cells to BYL719 treatments. 4T07 cells were treated either with BYL719 alone or in combination with VPS34IN1, and their proliferation was monitored over 72 hours. The combination significantly reduced the proliferation rate of 4T07, particularly at high doses of BYL719, resulting in a 30% decrease at 40 uM (Fig. S6E). We examined if VPS34IN1 could enhance the sensitivity of 4T07 cells to Pik3ca inhibition, and found that treating serum-starved 4T07 cells with BYL719 and VPS34IN1 significantly decreased mTORC1 activity in contrast to single drug treatments (Fig. S6F-G). However, VPS34IN1 did not enhance the BYL719 inhibition of mTORC1 activity after stimulating 4T07 cells with insulin (Fig. S6F-G), suggesting that Pik3c3 primarily regulates mTORC1 through amino acid signalling (35).

We questioned whether the differential dependence on and activity of Pik3c3 between the 4T1 and 4T07 could be attributed to specific genomic or genetic events. We examined the WGS data for the genomic status of the three PI3K classes. Besides the *Pik3ca* p.A423S missense mutation, the 4T07 cells harbored a *Pik3c3* missense mutation at a conserved residue (p.E562K) (Fig. S7A). This residue is located in a loop of the kinase domain of PIK3C3(43) (Fig. S7B-C) and has been observed mutated in two patients (E562A, malignant peripheral nerve sheath tumor; E562K, penile carcinoma, Supplementary Methods). However, the oncogenic potential of these mutations is unknown. In our analysis, the E562K mutation is predicted to have a moderate effect (Table S6). To determine whether this mutation increases PIK3C3 activity, we generated HeLa Flp-In T-REx cells expressing either wildtype or mutated PIK3C3 in a tetracycline inducible manner. Induction of both forms of PIK3C3 did not result in a significant difference in mTORC1 activity (Fig. S7D), suggesting that this mutation is unlikely to be responsible for the 4T07 cells’ mTORC1 hyperactivity. Overall, we did not identify any other notable genomic differences between the two lines that could explain the observed molecular and biological phenotypes (Fig. S7A).

Next, we examined whether the phenomena observed in the 4T1 and 4T07 models were also present in human BC. To do this, we examined early passages of patient-derived xenograft (PDX) derived BC cell lines for mTORC1 activity (Fig. S8A-B). This identified 2 potential PDX cell lines (1915 and 1735, (15)) with high mTORC1 activity. We focused on the PDX line 1915 because it originated from a patient with basal BC (15), similar to 4T1/4T07 cells, and because the associated patient experienced a metastatic relapse 1.5 years after tumor resection (Fig. S8C). WGS of the patient tumor and the PDX indicated the lack of *PIK3CA* activating hotspot mutations or *PTEN* deletion (like 4T1/4T07 cells), which are common causes of mTORC1 hyperactivity (Fig. S8C). PDX1915 had an amplification in the *PIK3CA* gene without corresponding overexpression in its transcript level compared to PDXs with WT or altered *PIK3CA* gene (Fig. S8D). We tested whether inhibiting PIK3C3 activity could reduce mTORC1 activity in PDX1915 cells and found that VPS34IN1 treatement decreased mTORC1 activity by approximately 50% (Fig. S8E-F). VPS34IN1 also significantly reduced the percentage of peripheral lysosomes in these cells (Fig. 5E-F). Furthermore, VPS34IN1 increased the percentage of cells with tubulated lysosomes (Fig. 5E and 5G), consistent with previous studies (44). To extend these findings, we screened three additional human BC cell lines, CAMA-1, T47D, and HCC70. These lines differ in their BC intrinsic subtype and dependency on *PIK3C3* (Fig. S8G). VPS34IN1 treatement reduced mTORC1 activity by approximately 50% in all three lines (Fig. S8H-I). Moreover, this treatment reduced the percentage of peripheral lysosomes in T47D and HCC70, but not in CAMA-1 cells (Fig. 5H-M).

In summary, inhibiting PIK3C3 activity in six murine and human BC cell lines reduced mTORC1 activity and shifted peripheral lysosomes to the perinuclear region in five of them. This supports that PIK3C3 is an upstream mediator of mTORC1 activity in BC, partially through lysosomal positioning.

### High mTORC1 or PIK3C3 activity in primary tumors correlates with worse clinical outcome

We next investigated whether mTORC1 activity could predict relapse incidence and/or poor clinical outcome in BC patients. Previous studies have shown that high expression of RPS6KB1(S6K1), correlates with worse distant metastasis-free survival in patients with early BC (45). However, expression levels alone may not fully reflect high mTORC1 activity. To address this, we used Gene Set Variation Analysis (GSVA) to generate an “mTORC1 score” for each tumor in the METABRIC and TCGA-BRCA cohorts, based on a previously annotated and curated mTORC1 signaling-mediated gene signature (46) (Fig. 6A). We observed that Basal and HER2-positive breast tumors exhibited higher mTORC1 activity scores compared to the less aggressive luminal subtypes (2) (Fig. 6B). Furthermore, patients with tumors characterized by a high mTORC1 signature score tended to have a poorer prognosis compared to those with low scores in the METABRIC dataset (*p<*0.0001) and with a trend towards a similar correlation in the TCGA dataset (*p=*0.1) (Fig. 6C). These results aligns with the established impact of mTORC1 activity in human cancer (47). To explore the clinical significance of PIK3C3 activity, we generated a PIK3C3-specific gene signature (see supplementary Methods). We found that Basal breast tumors had the highest PIK3C3 score in METABRIC and TCGA datasets (Fig. 6D). Similar to the mTORC1 signature, patients with high PIK3C3 scores had worse prognosis when compared to those with low PIK3C3 scores in the METABRIC (*p<*0.0001) and TCGA (*p*=0.022) datasets (Fig. 6E). These observations highlight the potential value of targeting the PIK3C3-mTORC1 singnaling axis in BC.

**Figure 6.**
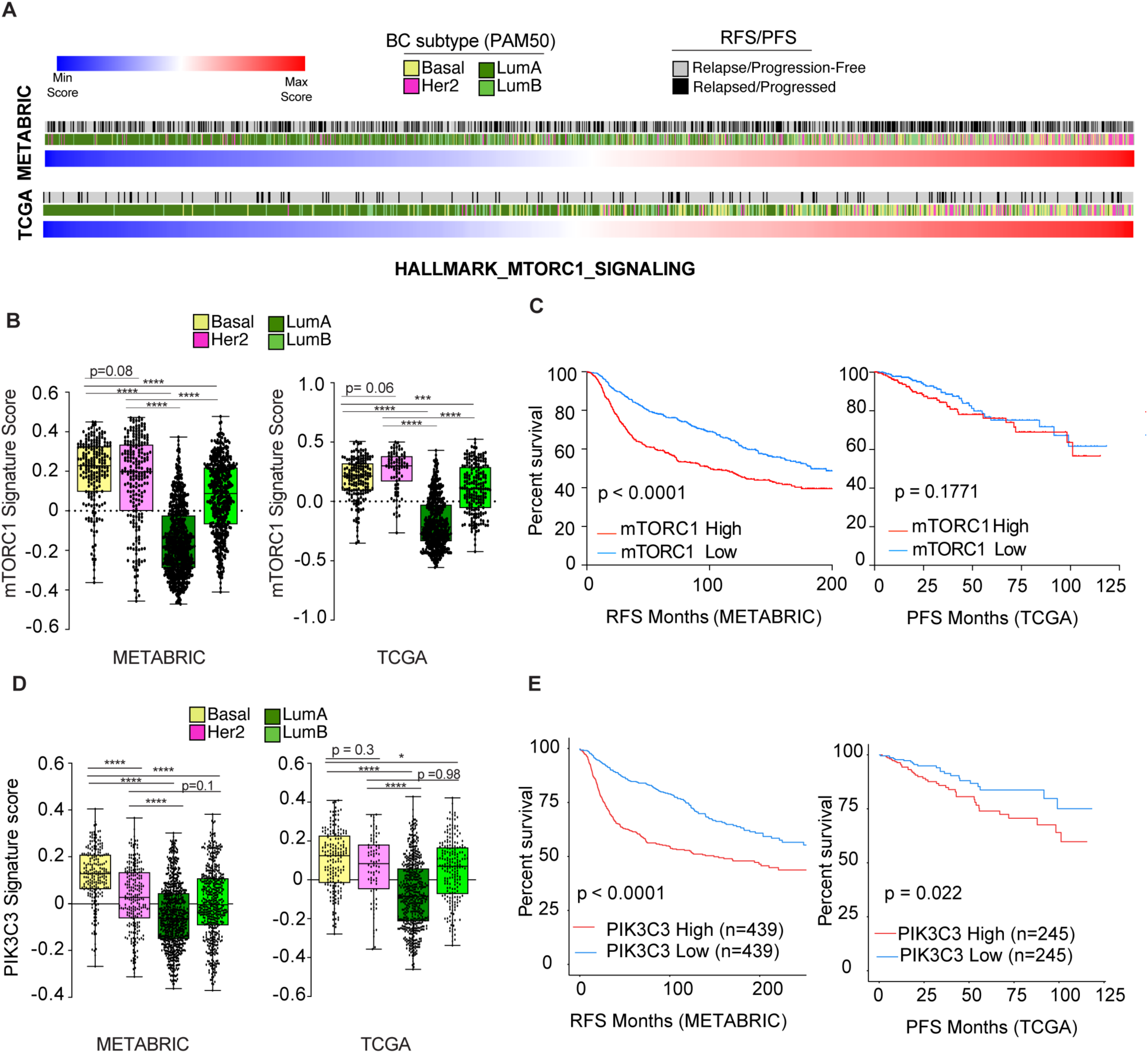
High mTORC1 or PIK3C3 activity correlates with worse outcome in breast cancer patients. **(A)** Heatmaps showing GSVA ranked by the Hallmark_mTORC1_Signaling signature score. Breast cancer samples were stratified according to their intrinsic molecular subtype based on gene expression profiling. **(B)** Comparing different breast cancer subtypes according to the Hallmark_mTORC1_Signaling signature score in the METABRIC and TCGA datasets. **(C)** Kaplan–Meier curves of disease-free survival and progression-free interval from the METABRIC and TCGA datasets, respectively. **(D)** Comparing different breast cancer subtypes according to the PIK3C3 signature score in the METABRIC and TCGA datasets. **(E)** Kaplan–Meier curves of Relapse-free survival and progression-free survival from the METABRIC and TCGA datasets, respectively.

### Inhibiting Pik3c3 decreases *in vivo* metastatic burden specifically in the 4T07 model

We explored whether targeting the PIK3C3-mTORC1 pathway in animal models could have a beneficial outcome on BC progression and metastasis burden. We performed spontaneous metastasis assays using the 4T07 and 4T1 models with or without treatement with VPS34IN1. 4T07 cells were implanted in the mammary fat pad of immunocompetent BALB/c mice, and primary tumors were allowed to grow for three weeks before the mice were randomely assigned to receive either vehicle or VPS34IN1 (Fig. 7A). After six days of treatment, the mice were sacrificed for analysis. There was no significant difference in primary tumor weight between the two groups, suggesting no effect on primary tumor burden (Fig. 7B). However, VPS34IN1-treated mice had approximately three-fold reduction in DTCs’ burden compared to vehicle-treated mice (Fig. 7C). To further investigate whether Pik3c3 inhibition could suppress the metastatic progression of the dormancy-prone cells when implanted under metastasis-permissive conditions, we implanted 4T07 cells expressing GFP and luciferase (4T07-TGL (12)) in the mammary fat pad of nude mice (Fig. S9A). One week post-implantation, the mice were randomized to receive either the vehicle or VPS34IN1 (Fig. S9A). Six days after treatment initiation, half of the mice were sacrificed for early timepoint analysis (Fig. S9A). There was no difference in tumor weight between the two groups (Fig. S9B). However, VPS34IN1-treated mice had significantly lower DTCs’ burden compared to the control group as assessed by biolumeniscence imaging (Fig. S9C-D). Further analysis was performed on the second group of mice after 12 days of treatment (3 weeks post implantation) with either VPS34IN1 or vehicle. While there was no statistically significant difference in tumor weight between the two experimental groups (Fig. S9B), the incidence of visible metastases in the VPS34IN1-treated group (1/5 mice) was 40% of that of the control group (3/6 mice) (Fig. S9E). Analyzing the digested lungs of mice that showed no visible lung metastases suggested that the VPS34IN1-treated mice demonstrated a trend towards lower DTC burden compared to controls (*p*=0.06) (Fig. S9F). Next, we asked whether VPS34IN1 affected mTORC1 activity in vivo. We found lower mTORC1 activity in the VPS34IN1-treated mice versus the control group (Fig. S9G-H). These data suggest that pharmacological inhibition of the Pik3c3-mTORC1 pathway decreases the DTCs’ burden, hence reducing the incidence of metastasis.

**Figure 7.**
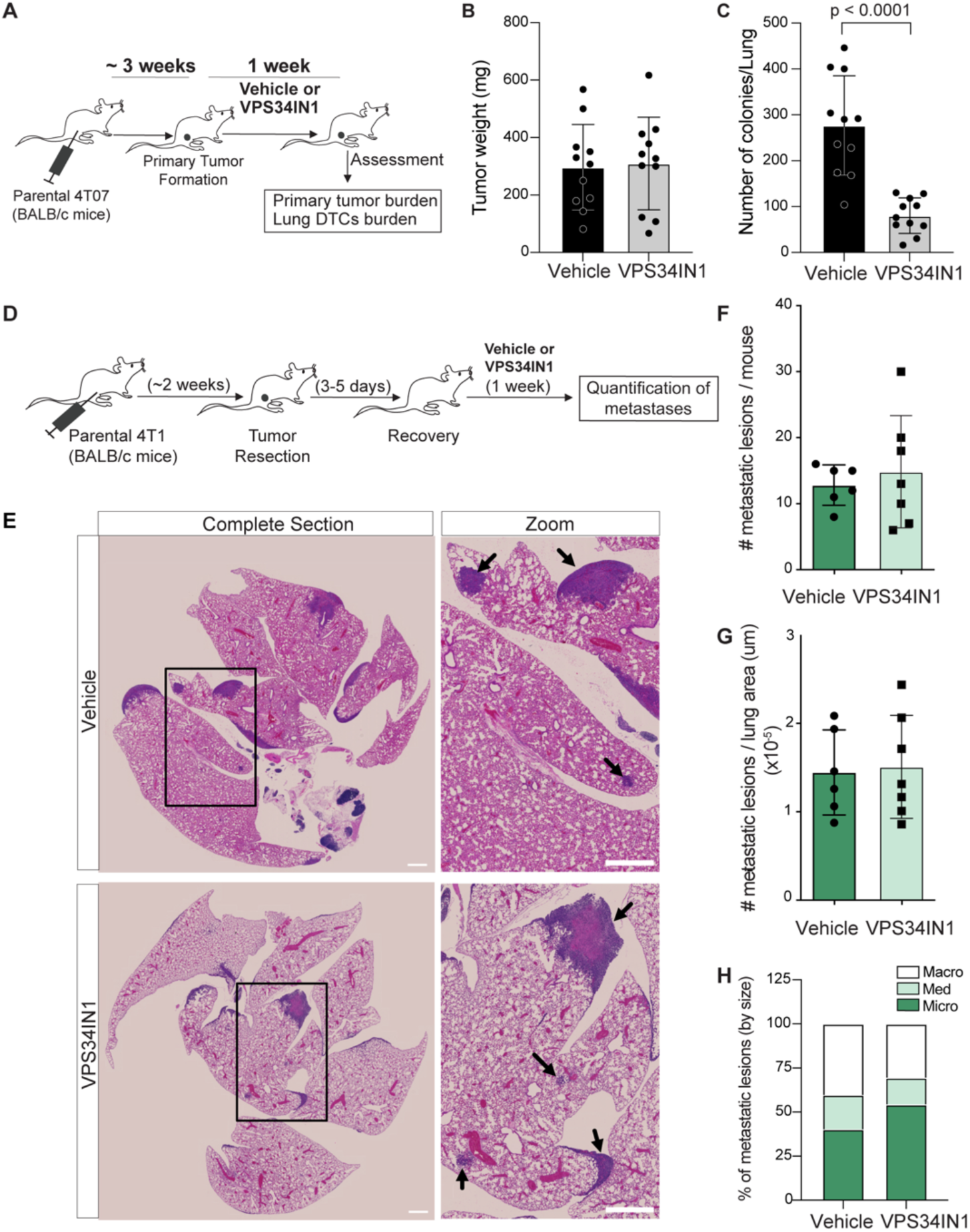
Inhibiting the Pik3c3-mTORC1 axis decreases the metastatic burden in vivo preferentially in the 4T07 model. **(A)** Schematic of the experiment investigating the effect of Pik3c3 inhibition on the metastasis burden in the 4T07 model. **(B)** Graph showing tumor weight of mice treated with either VPS34IN1 (50 mg/kg/day) or vehicle. **(C)** Graph showing the number of colonies retrieved from the lungs of mice treated with either VPS34IN1 (50 mg/kg/day) or vehicle. **(D)** Schematic of the experiment investigating the effect of Pik3c3 inhibition on the metastasis burden in 4T1 tumors-bearing BALB/c mice. **(E)** Representative H&E stainings of lungs obtained from mice in (D). Black arrows denote to metastatic lesions. **(F-H)** Quantifications of the metastasis burden: absolute number of lesions **(F)**, number of lesions normalized to lung area **(G)**, and percentage of different sizes of metastases **(H)** in the vehicle- and VPS34IN1-treated mice (50 mg/kg/day).

To investigate the role of Pik3c3 activity in metastatic progression in the 4T1 model, we implanted 4T1 cells in the mammary fat pads of immunocompetent BALB/c mice. Two weeks later, the primary tumors were surgically resected (Fig. 7D). After recovery, the mice were randomized to receive either VPS34IN1 or vehicle treatement for one week, followed by an assessment of lung metastasis burden via immunohistochemistry. There was no difference in the incidence of metastasis, as both experimental groups showed a 100% incidence rate (Fig. 7E). Moreover, there was no significant difference in the number or size of metastatic lesions (Fig. 7F-H). To maintain a consistent comparison between the 4T1 and 4T07 models, we implanted 4T1-TGL cells into the mammary fat pads of nude mice. After one week, the mice were treated with either VPS34IN1 or vehicle for two weeks before analyzing the primary tumor and metastasis burdens (Fig. S10A). There was no significant differences between the two groups in terms of primary tumor burden (Fig. S10B) or metastasis incidence (Fig. S10C), despite a trend towards reduced mTORC1 activity in the 4T1 primary tumors (Fig. S10D-E). Collectively, these results demonstrate that pharmacological inhibition of the Pik3c3-mTORC1 pathway can reduce BC metastasis, particularly in models with peripheral lysosomal positioning and high mTORC1 activity.

## DISCUSSION

The landscape of the cancer cells’ intrinsic factors contributing to their metastatic fate and response to therapies remains poorly outlined. Here, we compared two related basal B-like BC models, 4T1 and 4T07, which differ in their metastatic capabilities and dormancy potential. We show that metastatic cells exhibit higher PI3K activity and depend on it more than dormancy-prone cells. These observations align with earlier reports demonstrating that bone marrow-derived DTCs from BC patients during metastatic latency period show low PI3K activity (31,48). Notably, despite the lower PI3K-AKT activity observed in the dormancy-prone 4T07 cells (this study) and in D-HEp3 cells (model of dormant human head and neck carcinoma (31)), both cell lines exhibited higher mTORC1 activity compared to their metastatic counterparts, 4T1 and T-HEp3 (31), respectively.

A surprising finding in our study is that peripheral lysosomal positioning contributes to elevated mTORC1 activity in the dormancy-prone cells, despite their low PI3K activity. Previous studies have established that peripheral localization of lysosomes enhances mTORC1 activation, whereas perinuclear clustering reduces this activity and promotes macroautophagic dynamics (37,38). Our results suggest that the 4T1-4T07 cell line pair may follow this binary model. Exploring whether there is a differential autophagic potential between these two lines would be valuable in future studies given the conflicting reports on the role of autophagy in dormancy (21,49).

PIK3C3, activates mTORC1 (35) through various mechanisms (27,30,37,50), including promoting the anterograde transport of lysosomes to the cell periphery, where mTORC1 becomes activated (37). We found that inhibiting Pik3c3 activity reversed the persistent peripheral lysosomal positioning and the associated high mTORC1 activity in dormancy-prone cells. This inhibition also reduced the ability of these cells to colonize lung tissues, suggesting that this molecular machinery plays a role in successful metastasis. Consistent with a conserved role in human tumors, PIK3C3 inhibition mediated similar effects in three human BC lines with different genomic alterations and intrinsic subtypes. In one of the six murine and human cell lines we tested, CAMA-1, the reduction in mTORC1 activity was not accompanied by a shift in lysosomal positioning following PIK3C3 inhibition. This indicates that PIK3C3 may regulate mTORC1 activity though mechanisms independent of lysosomal positioning (30). Future research in this area could enhance our understanding of the genetic and signalling dependence of dormancy-prone cells.

mTORC1 and PIK3C3 have an entangled bilateral relationship. While we focused on PIK3C3’s role upstream of mTORC1, it has been reported that mTORC1 also acts upstream of PIK3C3 to mediate lysosomal regeneration following starvation-induced autophagy, supporting cell survival (44,51). These studies have identified tubulated lysosomes as transient autophagy-induced structures that serve as proto-lysosomes during lysosomal regeneration (51). However, these structures are also observed under nutrient-rich conditions (44), with their formation being dependent on mTORC1 activity. In the PDX1915 cells, we found that the presence of these structures was augmented upon treatment with VPS34IN1, despite the reduction in mTORC1 activity. It is unclear whether this increased tubulation is a feedback mechanism induced by reduced mTORC1 activity to maintain a basal lysosomal pool for mTORC1 activation. The potential connection between the lysosomal positioning, mTORC1 activity, autophagy, and the formation of lysosomal tubules warrants further investigation.

Our findings provide insights into research directions of potential clinical relevance (52). Gene signatures associated with either mTORC1 or PIK3C3 correlated with clinical outcome in BC. However, when stratifying patients by intrinsic BC subtype, no statistically significant correlation was observed with either signature in TCGA or METABRIC datasets. One exception is a significant correlation between the PIK3C3 signature and outcome in patients with Luminal A breast cancer (not shown). This might be due to the sample size, suggesting that analyzing larger cohorts with different subtypes could provide more insights. It is important to note that the PIK3C3 signature used was generated from datasets from pancreatic and lung cancers. Given the relatively unexplored roles of PIK3C3 in BC pathogenesis, future studies aiming at generating BC-specific PIK3C3 signatures might be of significance. From a therapeutic perspective, Gedatolisib, a dual Class-I PI3K/mTOR inhibitor, failed to reduce the metastatic burden in two BC models when used alone or combined with standard-of-care chemotherapy (53). Another study demonstrated that long term pharmacological inhibition of PI3K or AKT induces a PIK3C3-SGK3 signaling axis to activate mTORC1, and targeting this axis can revert the induced resistance towards the PI3K-AKT inhibition (30). Our work demonstrating that dormancy-prone cells do not depend on the class-I PI3K and are resistant to its pharmacological inhibition meanwhile being dependent on the class-III PI3K complements these studies. Future studies should explore whether the combinatory inhibition of class-I and -III PI3K, and mTORC1 could effectively reduce the metastatic burden in vivo. Importantly, the contribution of microenvironment to the observed effect of inhibiting PIK3C3 on BC metastasis should be considered.

While our CRISPR screening approach revealed novel insights into the intrinsic differences between cells of differential metastatic capacity, it has limitations. The screens were performed for technical feasibility in an *in vitro* setting that does not necessarily recapitulate the impact of the cross-talk with microenvironment on gene essentiality and signaling pathways’ activity. Nonetheless, we extensively validated the PI3K(s)-mTORC1 signaling circuits in vitro and in vivo. Another limitation of our study is depending on one pair of cell lines with validations in a patient-derived xenograft and three human cell lines. Our molecular findings are reaffirmed in these additional models. Whether the relevance of these findings in the context of dormancy could be extended in other models of different genomic backgrounds would be worth addressing in future studies. Such a limitation goes back-to-back with the global challenge of scarce authentic cell line pairs that possess different metastatic potentials upon being implanted in syngeneic mice (8).

In summary, the work described here provides novel insights into the intrinsic differences between cells of differential metastatic capacity that can guide future explorations of the biology and therapeutic vulnerabilities of BC metastatic relapses.

## ACKNOWLEDGMENTS

We would like to thank Dr. Marie-Anne Goyette for providing constructive feedback on the manuscript. We acknowledge Dr. Flippo Giancotti (4T07-TGL cell line) and Dr. Vladimir Ponomarev (TGL reporter) for providing reagents. We thank Drs. Sandra Turcotte, Philippe Roux, Matthew Smith, Anne-Marie Fortier and Deepak Singh for fruitful discussions. We acknowledge the “Genome Engineering using CRISPR/Cas Systems” Google discussion group, Dr. Julia Joung, and Dr. Jonathan Boulais for helpful discussions on technical steps or the bioinformatic analysis of the CRISPR screens. We express our gratitude to the IRCM facilities staff (Dr. Dominic Filion, Manon Laprise, and Dr. Odile Neyret) for their expert technical help involved in this work.

This work was supported by operating grants from the Cancer Research Society (operating grant #25244 to J.-F.C), Canadian Institutes of Health Research (CIHR) (Foundation grant #FDN-143281 to M.P. and operating grant #PJT-156086 to C.L.K), Canadian Cancer Society and Oncopole grants (to M.P.) and an American Cancer Society Research Scholar Award (RSG-22-164-01-MM to A.P.G.). PDX cell lines were obtained from the breast tissue bank at McGill University, supported by grants to the Reseau Cancer banque de tumeurs from the Fonds de la Recherche du Québec-en Santé (FRQS) and the Quebec Breast Cancer Foundation (to M.P). Data computational analyses were enabled by compute and storage resources provided by Compute Canada and Calcul Québec. I.E.E. was a recipient of an FRQS Doctoral Scholarship and an IRCM Foundation-TD scholarship. I.E.E. is currently supported by Peter Quinlan Postdoctoral Research Fellowship in Oncology (McGill University) and an FRQS Postdoctoral Fellowship. S.D. is a recipient of a Miles for Moffitt postdoctoral fellowship. C.L.K. is the recipient of an FRQS Salary Award. J.A.A-G. is supported by the following awards: National Institute of Health (NIH) /National Cancer Institute (NCI) (CA109182, CA253977, CA284085, P30CA013330), The Mark Foundation Aspire program, The Gurwin Foundation, DoD-MRP, DoD-BCRP, Melanoma Research Alliance and the Rose C. Falkenstein Chair in Cancer Research. J.A.A-G. is a Samuel Waxman Cancer research Foundation Investigator. M.P. is a Distinguished James McGill Professor and holds the Diane and Sal Guerrera Chair in Cancer Genetics. J.-F.C. holds the Canada Research Chair Tier 1 in Signalling in Cancer and Metastasis and the Alain Fontaine Chair in Cancer Research from the IRCM Foundation.

## AUTHOR CONTRIBUTIONS

Conceptualization, Supervision, Writing-Finalizing the manuscript: IEE and J-FC. Methodology & Investigation: IEE, AR, CM, HK, SD, GM, SH, AP, VC. Formal analysis & Writing - Review & Editing: all authors. Writing—original draft: IEE

## DATA AND MATERIALS AVAILABILITY

All data are available in the main text or the supplementary materials. RNA-Seq related datasets/analyses (e.g., 4T1-4T07 differentially expressed genes and correlation with human breast cancer cell lines) are included in tables S1 and S8. Moreover, the raw data of the RNA-Seq dataset is deposited online (GSE203296) and have been publicly available since December 1st, 2023. CRISPR screens’ data are provided in tables S2-5. WGS datasets are provided in tables S6-7.

## Notes

**Conflict of Interest Disclosure** Julio A. Aguirre-Ghiso is a scientific co-founder, scientific advisory board member and equity owner in HiberCell and receives financial compensation as a consultant for HiberCell, a Mount Sinai spin-off company focused on the research and development of therapeutics that prevent or delay the recurrence of cancer. The remaining authors declare no competing interests.

### Competing Interest Statement

COMPETING INTEREST:
Julio A. Aguirre-Ghiso is a scientific co-founder, scientific advisory board member and equity owner in HiberCell and receives financial compensation as a consultant for HiberCell, a Mount Sinai spin-off company focused on the research and development of therapeutics that prevent or delay the recurrence of cancer. The remaining authors declare no competing interests.

### Summary of Updates

This is a revised version of the manuscript that addresses reviewers' comments through the peer review process.

