## Supplemental data and methods for "Targeting the dependence on PIK3C3-mTORC1 signaling in dormancy-prone breast cancer cells blunts metastasis initiation"

#### SUPPLEMENTARY MATERIAL

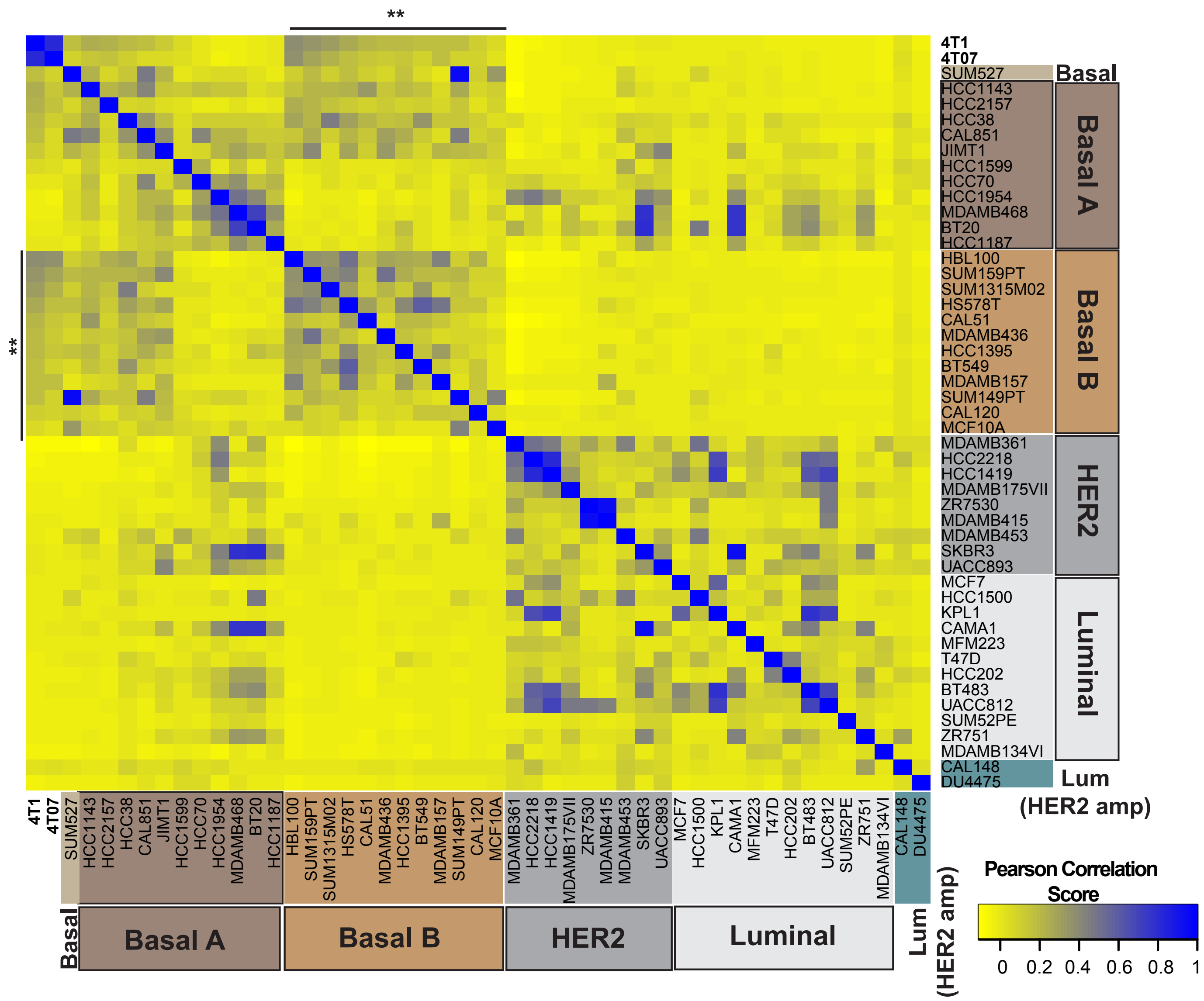


**Figure S1. Correlating the 4T1 and 4T07 transcriptomic profile to 47 human breast cancer cell lines.** Heatmap showing that the 4T1 and 4T07 cells scored the highest Pearson correlation score with the basal (basal-like) B breast cancer cell lines.

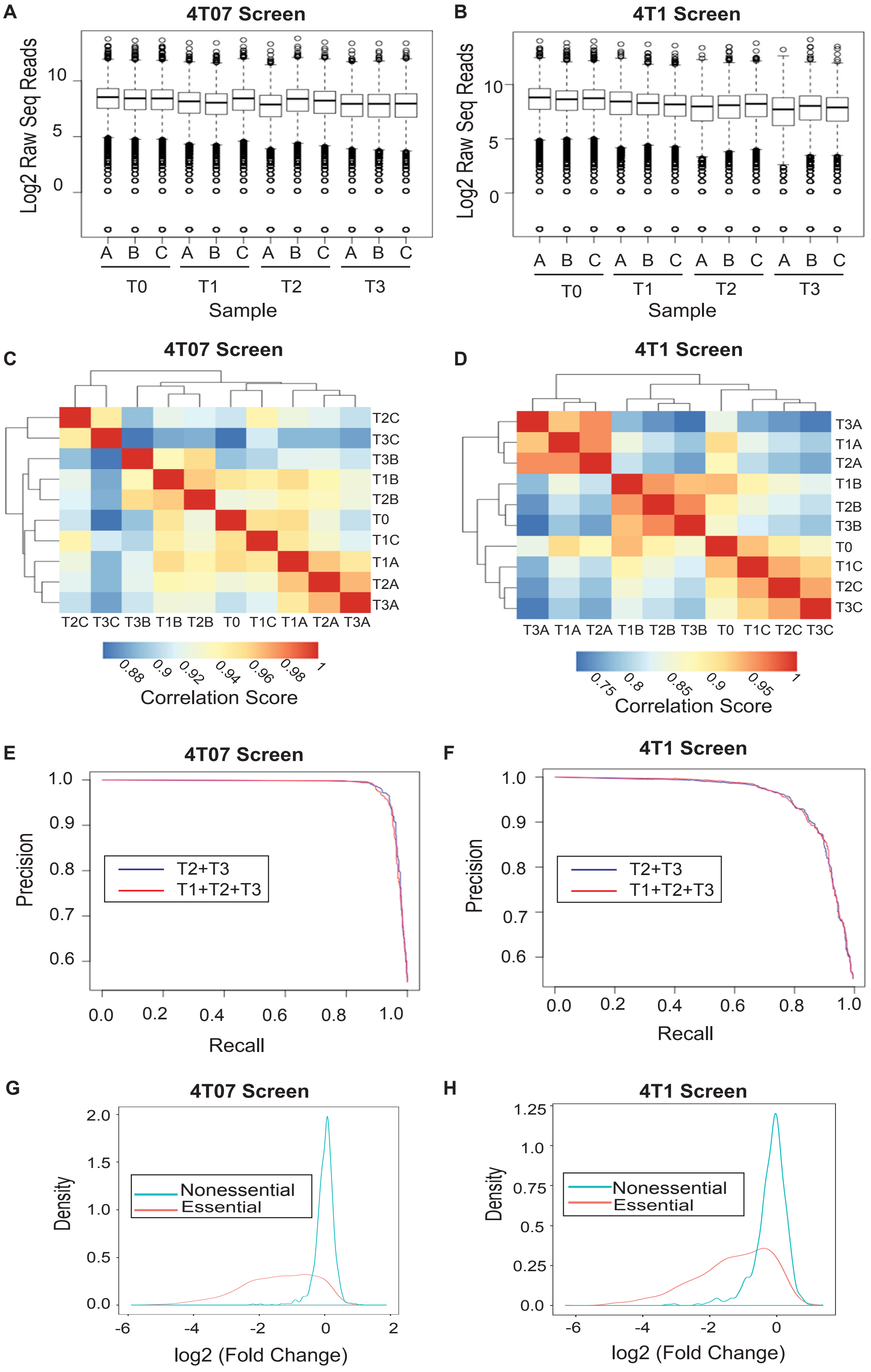


**Figure S2. Technical validations of the CRISPR screens. (A-B)** Boxplots showing the raw sequencing reads of the two screens. Each timepoint (T0, T1, T2, and T3) was replicated biologically three times (A, B, and C). **(C-D)** Heatmaps assessing the correlation between the different replicates and timepoints. **(E-F)** Precision-Recall curves assessing the ability of the performed screens to recall the previously annotated fitness genes. The results did not change when either the combination of T2 and T3 or the combination of T1, T2, and T3 was used for the analysis. **(G-H)** Fold-change distribution of gRNAs targeting previously annotated fitness genes (orange) or non-essential genes (blue) showing gRNAs targeting fitness genes are depleted as expected.

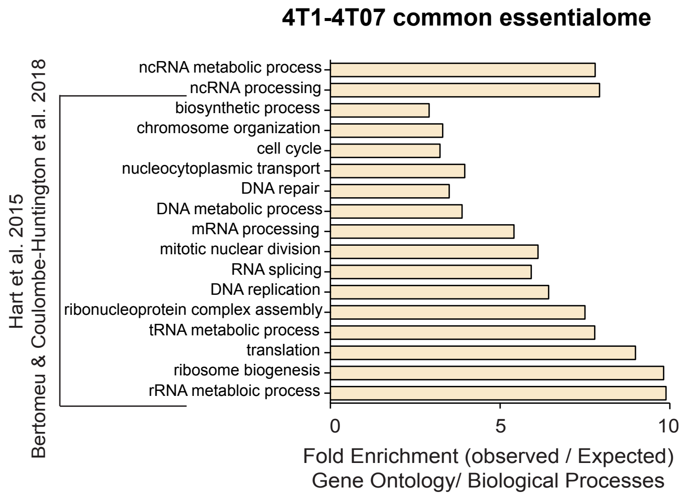


**Figure S3. Functional analysis of the 4T1 and 4T07 common fitness genes.** Enriched biological processes for the predicted common essential 4T1-4T07 genes including multiple processes previously described for the core fitness genes shared between different cancer cell lines (13,25).

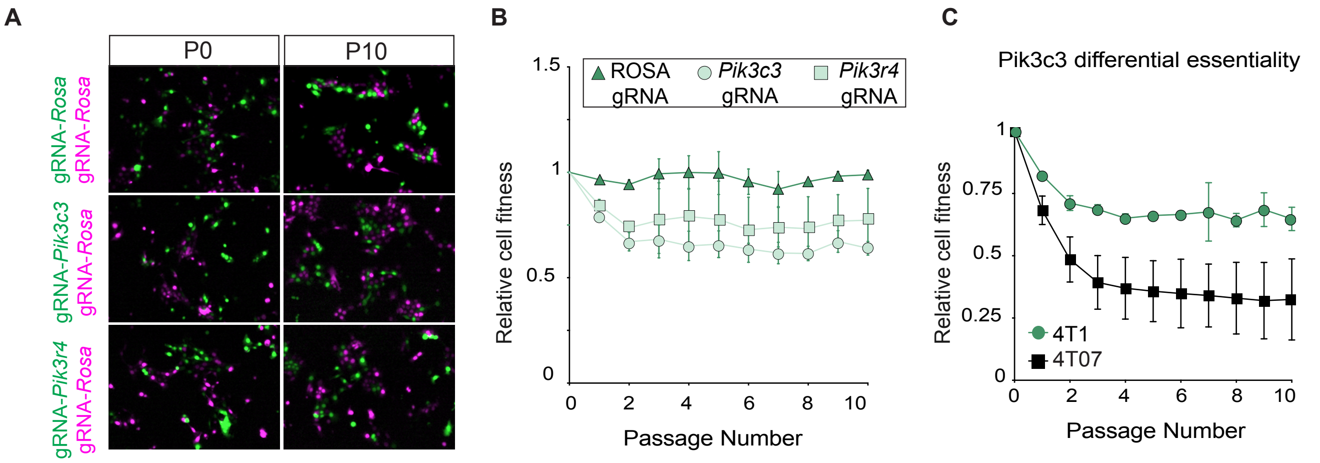


**Figure S4. Differential essentiality of *Pik3c3* in the 4T1 and 4T07 cells. (A)** Representative images showing either 4T1 cells expressing *Pik3c3*-gRNA, *Pik3r4*-gRNA, or *Rosa26*-gRNA (all in green) co-cultured independently with cells expressing *Rosa26*-gRNA in magenta, at passages zero (P0) and ten (P10). **(B)** Graph showing representation of the green 4T1 cell populations over serial passages in reference to P0 (n=2). **(C)** Direct comparison for the representation of the gRNA-*Pik3c3* conditions in (B; 4T1 cells) and (Figure 2G; 4T07 cells) after normalizing to the respective gRNA-*Rosa26* condition.

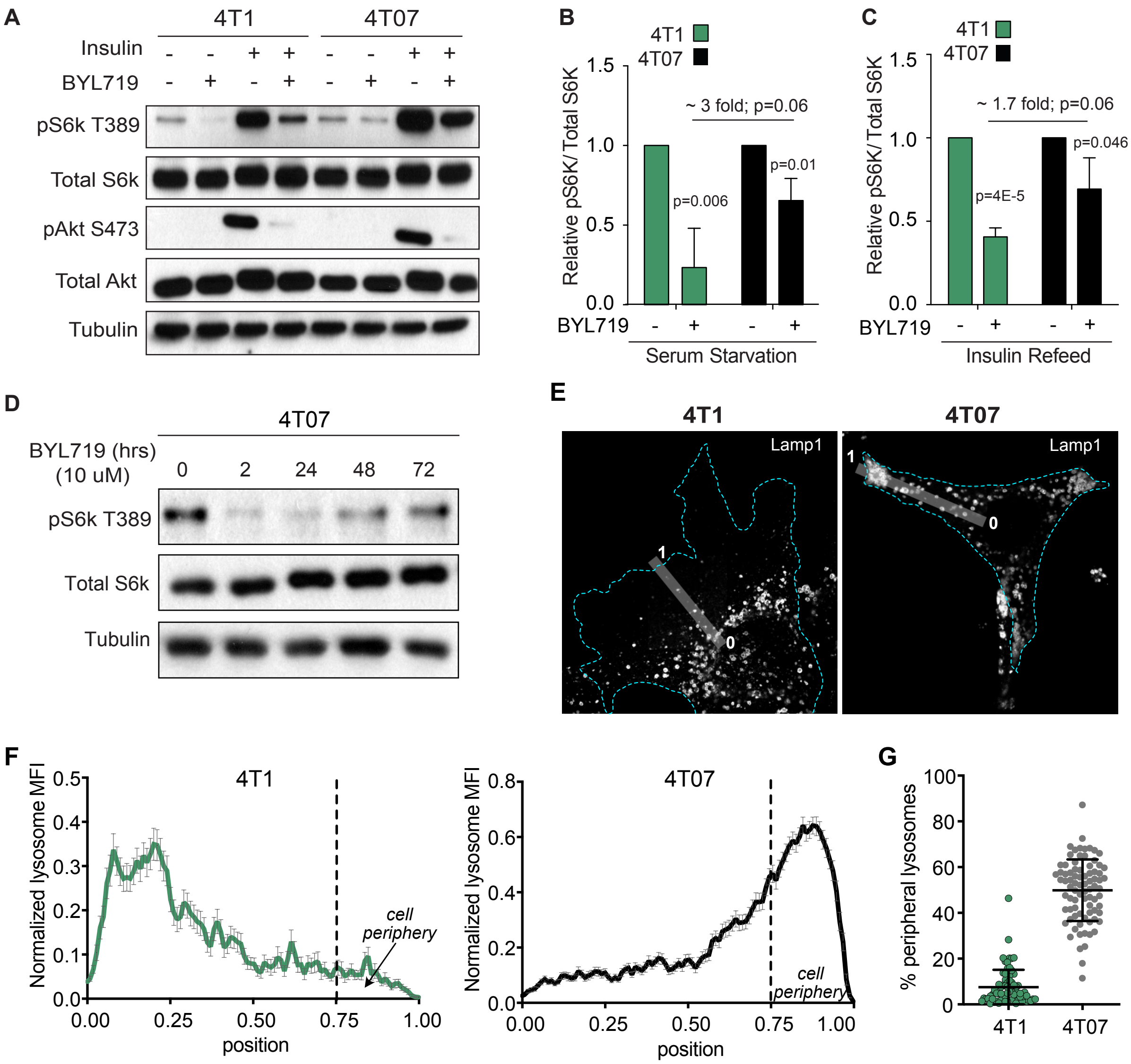


**Figure S5. Differential mTORC1 activity and lysosomal positioning between the 4T1 and 4T07 cells. (A)** Western blot analysis assessing the Pik3ca activity inhibition (1 µM) in the two cell lines under serum starvation and insulin refeeding. **(B-C)** Quantification of the mTORC1 activity investigated in panel (A). **(D)** Western blot analysis of mTORC1 activity at different timepoints under the effect of BYL719. **(E-G)** Plot profile analysis of lysosome distribution. The illustrations show an example of 4T1 and 4T07 images used for quantifications. The green channel (i.e., pMtor labeling as in Figure 4) was used to trace a 5 pixel-wide line from position 0 (border of the nucleus) and 1 (cell edge). The intensity profile for the red channel (Lamp1 staining) along that line was obtained using the plot profile plugin in FIJI. The results were normalized using MatLab and graphs in (F) represent the normalized data (Mean +/- SEM) from at least 75 cells per condition from 3 independent experiments. The dashed line represents the three quarters of the line that was used to separate the element present at the cell periphery (between 0.75 to 1). These graphs were used to calculate the area under the curve at the cell periphery versus total presented in panel (G) (Mean +/- SD). The same method was used to quantify the distribution of lysosomes presented in (Fig. 4D, 5D, 5F, 5I, 5K, and 5M). **(G)** Representative summary of the quantifications in (F).

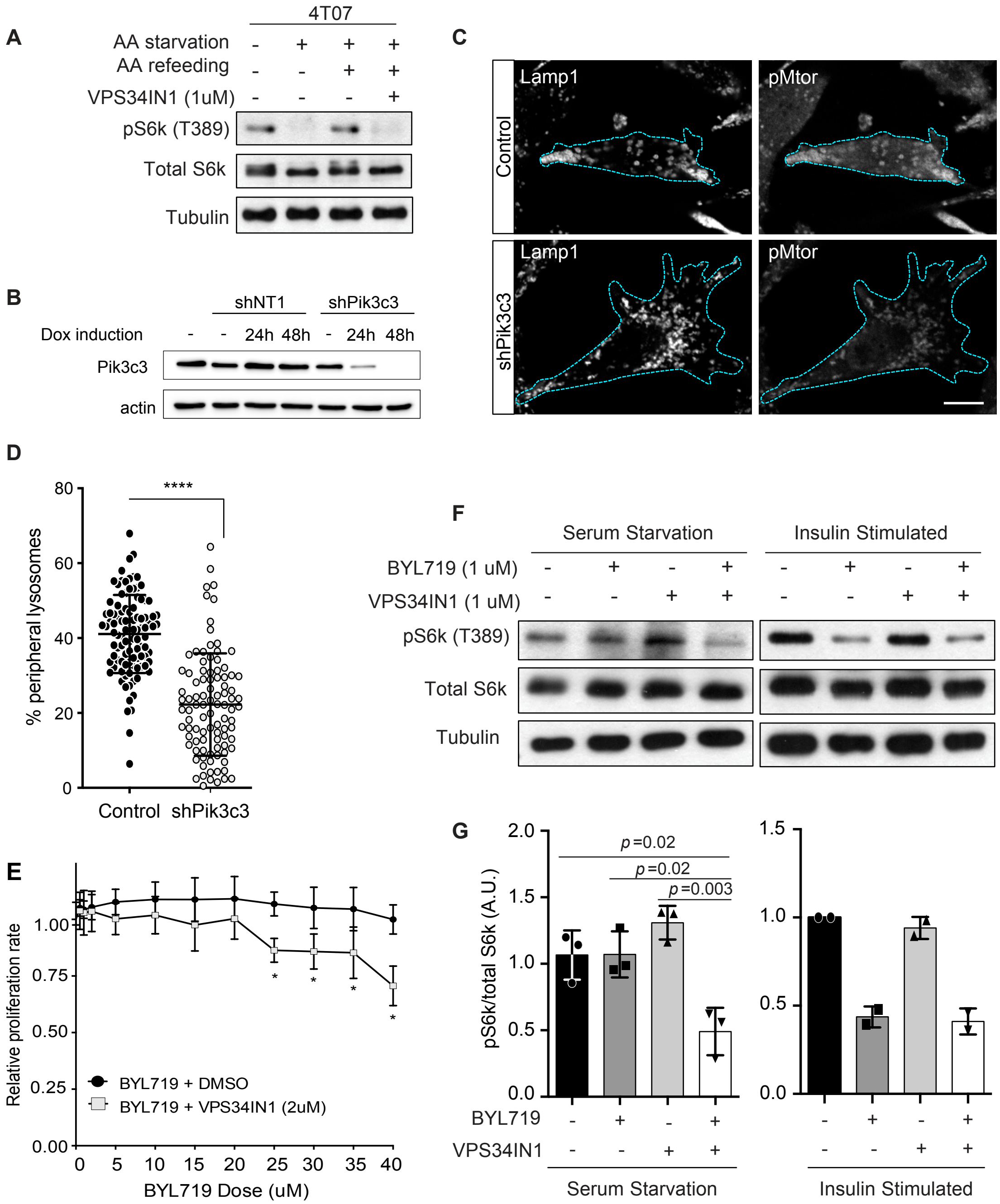


**Figure S6. VPS34IN1 sensitizes the 4T07 cells to Pi3k inhibition by BYL719. (A)** Western blot analysis for the effect of VPS34IN1 on mTORC1 activity under amino acid (AA) starvation and refeeding. **(B)** Validation of the 4T07 doxycycline-inducible *Pik3c3* shRNA cell line by western blot. The expression of non-Targeting shRNA control (shNT1) and Pik3c3 shRNA were induced with 1μg/mL doxycycline for the indicated time. **(C)** Immunostaining for Lamp1 and pMtor in 4T07 shPik3c3 non-induced (control) or induced for 48h with doxycycline. **(D)** Quantifications of the percentage of peripheral lysosomes in 4T07 cells 48 hours after inducing the Pik3c3 shRNA in comparison to the control condition (non-induced). **(E)** Proliferation assay comparing 4T07 cells' response to BYL719 alone or in combination with VPS34IN1 (*$p$= 0.009 at 40 μM). **(F-G)** Western blot analysis for the effect of combined inhibition of Pik3ca and Pik3c3 activity in 4T07 cells under serum starvation and insulin refeeding.

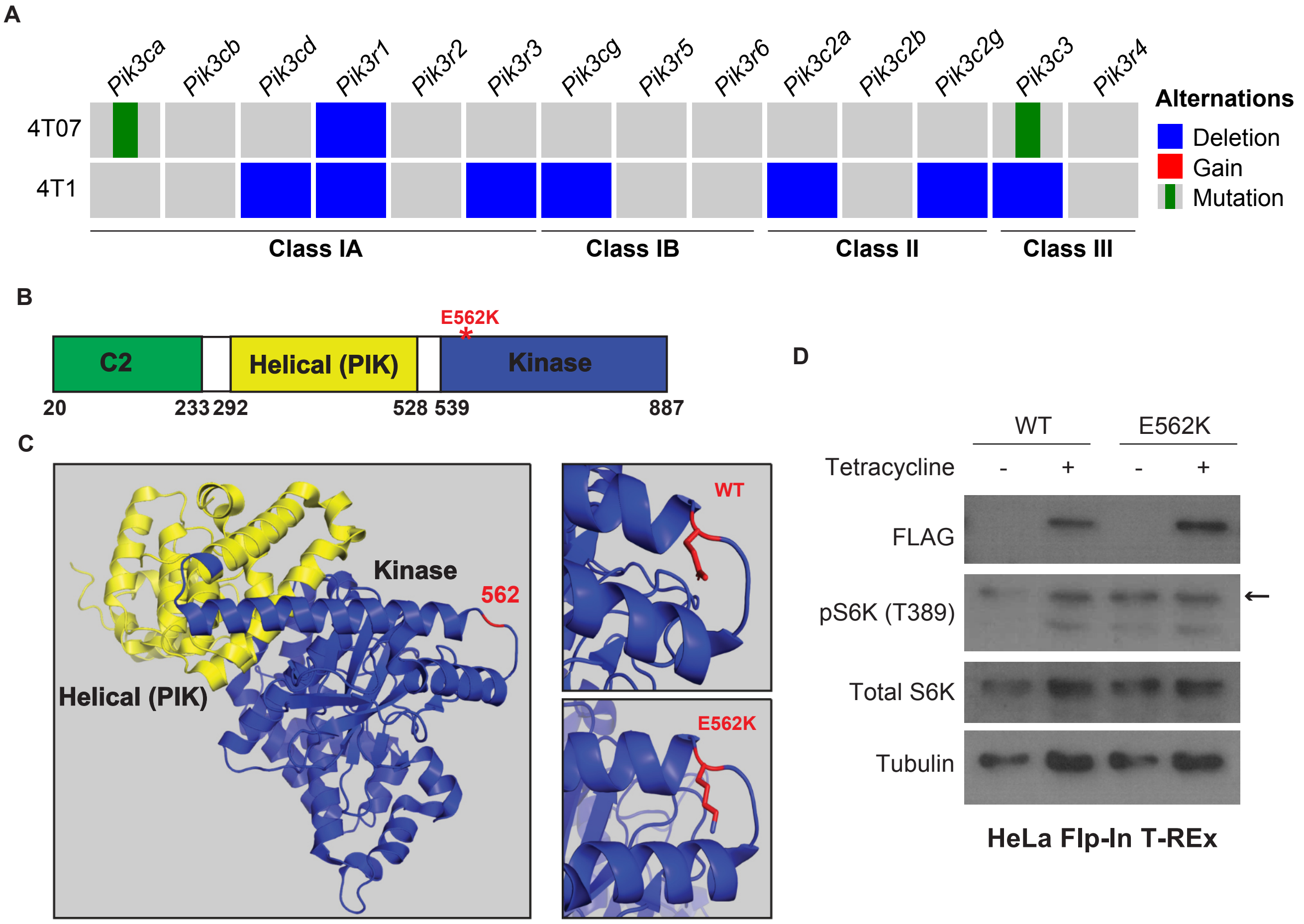


**Figure S7. Investigating the effect of PIK3C3 p.E562K mutation on mTORC1 activity**. **(A)** Oncoprints for somatic alterations in genes encoding different classes of Pi3ks (catalytic and regulatory subunits). **(B)** Schematic of the PIK3C3 protein's domains, highlighting the E562K mutation in the kinase domain. **(C)** Representation of the PIK3C3 structure and the location of the E562K mutation. WT; wild type. **(D)** Western blot analysis of mTORC1 activity after inducing the expression of WT or mutant (E562K) PIK3C3 in HeLa cells.

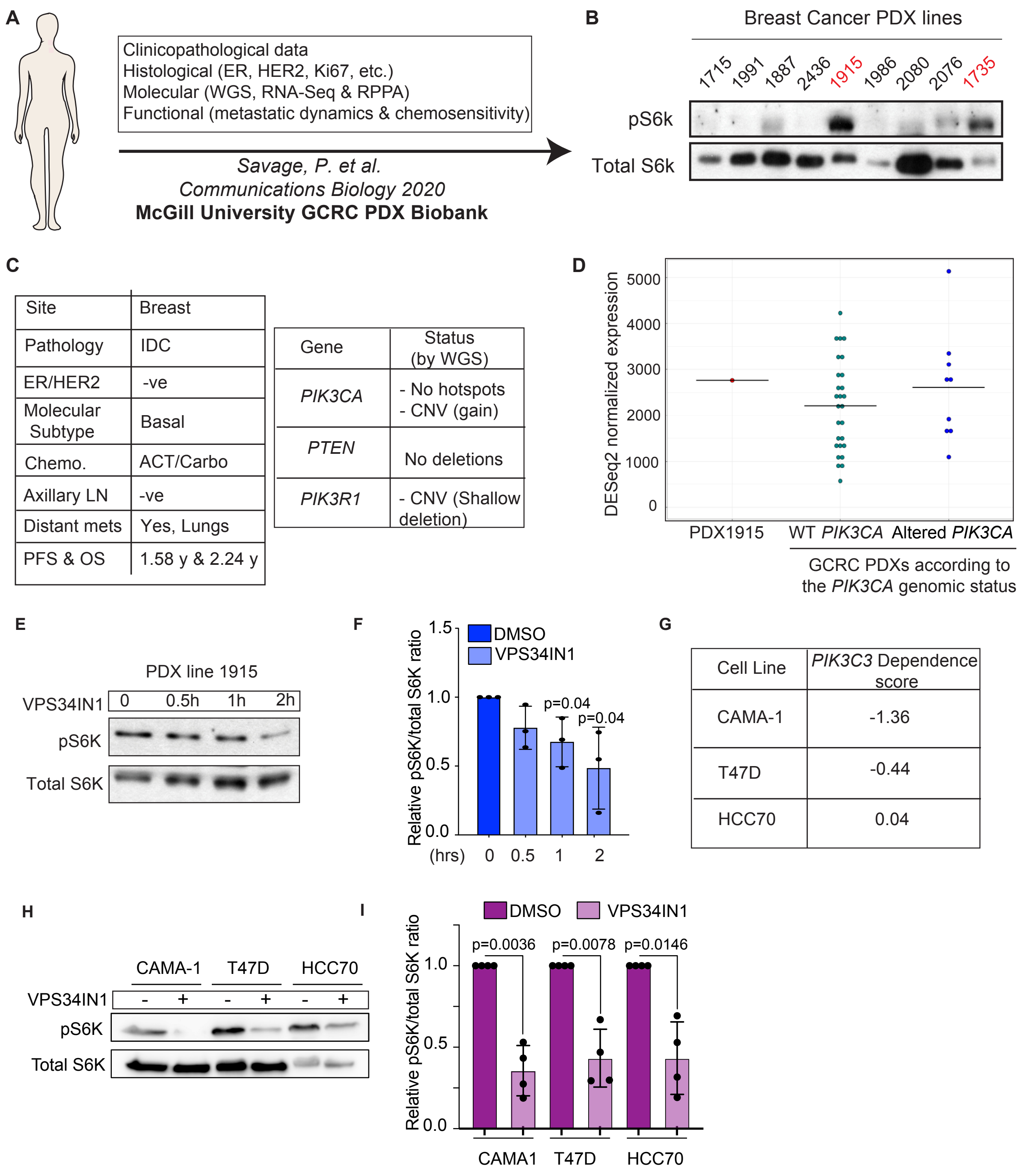


**Figure S8. Investigating the mTORC1 activity in the context of human breast cancer metastatic relapse. (A)** Schematic for the McGill University GCRC Patient-Derived Xenograft (PDX) biobank (15). **(B)** Western blot analysis of mTORC1 activity in 9 PDX lines. **(C)** Tables summarizing the clinicopathological characteristics of patient from whom the PDX 1915 originated in addition to the mutational status of the main genes in the PI3K pathway as assessed by whole genome sequencing (WGS). IDC, invasive ductal carcinoma; chemo, chemotherapy; ACT, Adriamycin-Cyclophosphamide-Taxol regimen; carbo, Carboplatin; LN, lymph node; mets, metastasis; PFS, progression-free survival; OS, overall survival. **(D)** Graph showing the mRNA expression level of *PIK3CA* in the PDX1915 in comparison to the rest of the PDXs in the biobank. **(E)** Western blot analysis for the VPS34IN1 effect on mTORC1 activity in the PDX-1915 (1 μM for 2 hours). **(F)** Quantification for the mTORC1 activity in panel (E). **(G)** Table summarizing the PIK3C3 dependence score in CAMA-1, T47D, and HCC70 cells as predicted by the DepMap portal. Cell lines are ranked from the highest dependence score to the lowest. Note that negative score indicates higher dependence on *PIK3C3*. **(H)** Western blot analysis for the VPS34IN1 effect on mTORC1 activity in the CAMA-1, T47D, and HCC70 cells (2 μM for 2 hours). **(I)** Quantification for the mTORC1 activity in panel (H).

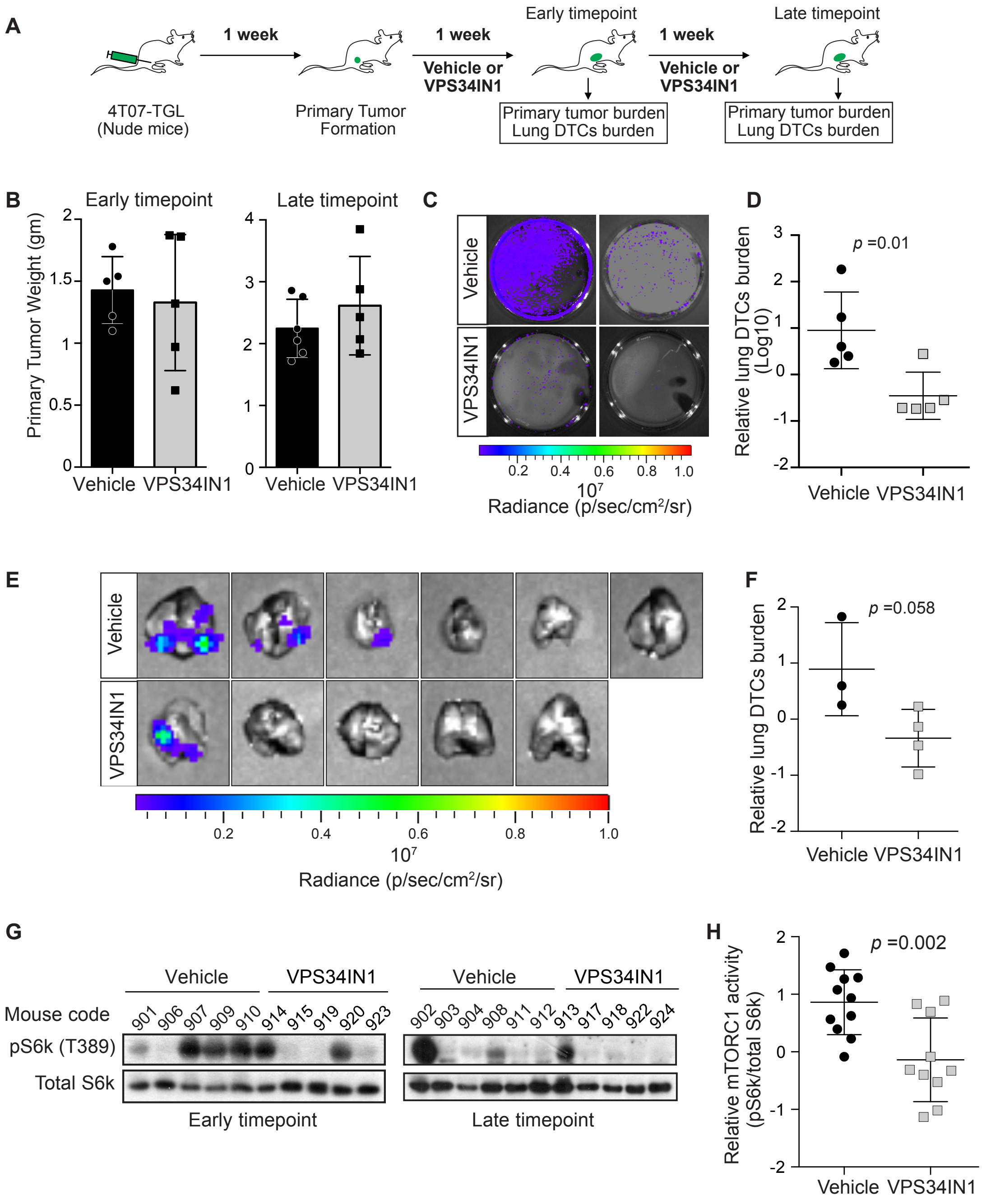


**Figure S9. Targeting the Pik3c3-mTORC1 axis decreases the in vivo metastatic burden in the 4T07 model. (A)** Schematic of the experiment investigating the effect of Pik3c3 inhibition on the metastasis burden of the 4T07 model in nude mice. **(B)** Graph showing tumor weight of mice treated with either VPS34IN1 (50 mg/kg/day) or vehicle at the early and late experimental endpoints. **(C)** Representative images of the lung DTCs burden in 4T07-bearing mice treated with VPS34IN1 or vehicle at the early experimental endpoint (n=5/group). **(D)** Quantification of relative lung DTCs burden between the two groups in (C). **(E)** Representative ex-vivo lung images of 4T07-bearing mice treated with VPS34IN1 or vehicle at the experimental late endpoint. **(F)** Quantification of relative burden of the lung DTCs in mice with no visible metastases at the late experimental endpoint (from lungs presented in panel E). **(G)** Western blot analysis for mTORC1 activity in the 4T07 tumors from mice treated with either the vehicle or VPS34IN1 at the early or late experimental endpoints. **(H)** Quantification of the relative mTORC1 activity between the two groups in (G).

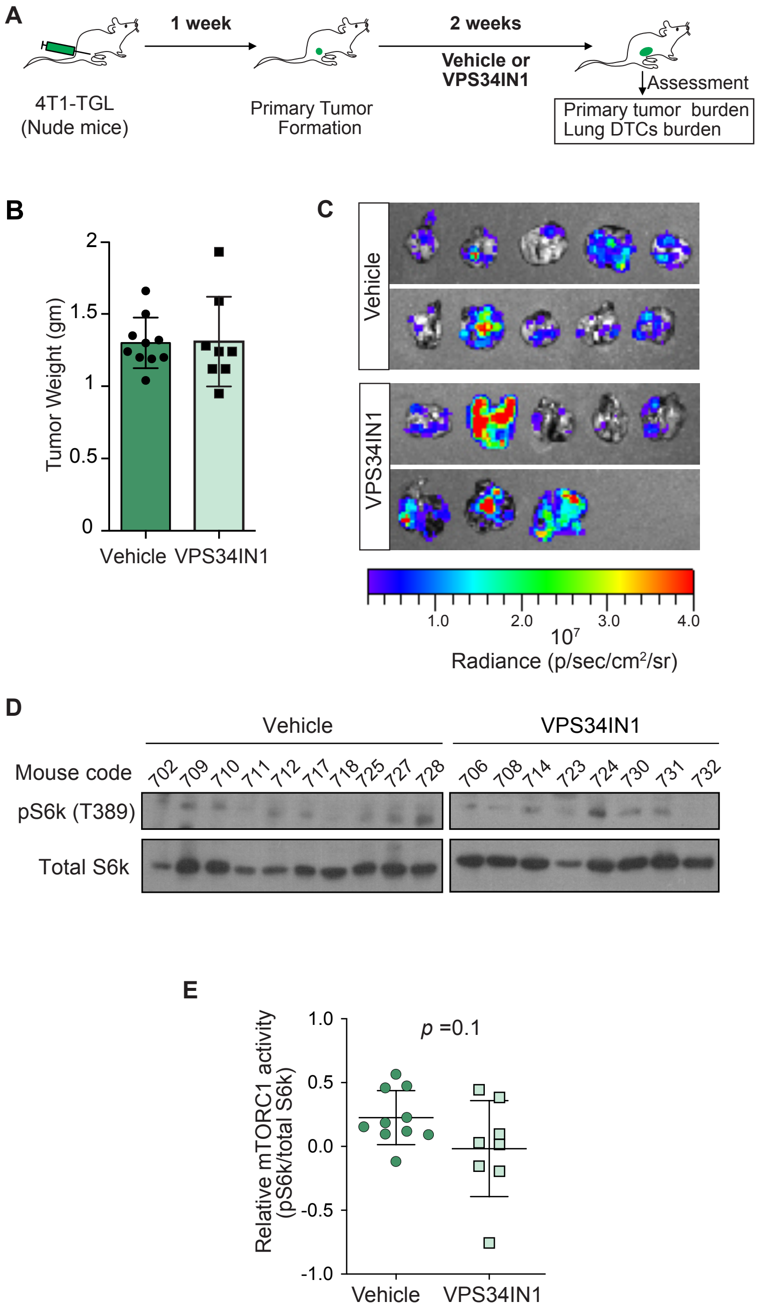


**Figure S10. Targeting the Pik3c3-mTORC1 axis does not affect the in vivo metastatic burden in the 4T1 model. (A)** Schematic of the experiment investigating the effect of Pik3c3 inhibition on the metastasis burden in 4T1 tumors-bearing nude mice. (**B)** Graph showing tumor weight of mice treated with either VPS34IN1 (50 mg/kg/day) or vehicle at the experimental endpoint. **(C)** Representative ex-vivo lung images of 4T1 tumors-bearing mice treated with VPS34IN1 or vehicle at the experimental endpoint. **(D)** Western blot analysis for mTORC1 activity in the 4T1 tumors from mice treated with either the vehicle or VPS34IN1. **(E)** Quantification of the relative mTORC1 activity between the two groups in (D).

### SUPPLEMENTARY METHODS

#### CRISPR-Cas9 Screen Analysis

The "count" and "test" commands of MAGeCK (1) (Version 0.5.7) were used to generate read counts for each gRNA and to identify positively selected genes in T3 compared to T0, respectively. Sequencing defined a range of 483-744 fold library coverage for the 4T07 screen and 529-881 fold library coverage for the 4T1 screen, significantly exceeding the recommended 100-300 fold in coverage. The BAGEL software (2,3) (v0.91) was then used to calculate the foldchange of gRNA representations in different timepoints in reference to T0. Consequently, BAGEL assigned each gene a Bayes Factor (BF) as a measure of confidence of fitness or essentiality, based on a pre-set list of essential and non-essential genes (2,4,5). This analysis was performed separately for each timepoint and using the final 2 timepoints combined, as previously described (6). For downstream analysis, we utilized BF scores from the final 2 timepoint analysis. A strict threshold of normalized BF > 6 was used to consider a gene essential in our screens (5). The generated BF scores were then used for the Precision-Recall analyses based on previously annotated gene lists (essential and non-essential) (results are found in Figure S2E-F). To compare the essential genes' profile of the two cell lines, the difference between the BF scores of the two lines was calculated. Then, a Z-score for the difference in BF for each gene was calculated (Table S2-4). Positive Z-score denoted 4T1-specific essential gene and negative Z-score denoted 4T07-specific essential gene.

#### RNA-Seq, cell line correlation analyses, and mutational analysis

4T1 and 4T07 cells were grown in 6 well plates. Total RNA was extracted and purified using the Qiagen RNeasy column kit (#74104) following to the manufacturer's instructions. RNA quality was assessed on an Agilent 2100 bioanalyzer, ensuring a RIN above 8.3. For libraries preparation, 1ug of total RNA was processed using the NEBNext Poly(A) mRNA Magnetic Isolation Module (New England Biolabs) and the KAPA stranded RNA-Seq library kit (Roche Diagnotics). The size distribution of the libraries was assessed using Agilent 2100 bioanalyzer. Libraries were loaded equimolarly on a lane of HiSeq 2500 and sequences at the McGill University and Génome Québec Innovation Centre (MUGQIC). After quality control with FASTQC (7), mapping was performed with Star v2.5 on GRCm38 reference genome (8). Around 85% of the reads were uniquely mapped. Transcripts were quantified with featureCounts v1.4.6 (9) using Ensembl v84 reference annotation and used as input for differential expression analysis with the DESeq2 R package (10). The generated dataset has been deposited with a GEO accession number: GSE203296. Functional analyses of the differentially expressed genes between 4T1-4T07 were performed using the Gene Ontology Resource (11,12) and Gene Set Enrichment Analysis (GSEA V3) tools (13,14).

To correlate the transcriptomic profile of 4T1 and 4T07 cells with human breast cancer cell lines, FPKM values for 675 human cancer cell lines (15) were extracted from https://www.ebi.ac.uk/arrayexpress/experiments/E-MTAB-2706/ and imported into R. Orthologs between human and mouse were extracted using Biomart (16). Data from human and mouse cell lines were merged based on orthologs. Initial analysis included 61 human breast cell lines. However, only 47 cell lines with well-defined subtyping (as identified in the depmap portal(17) and Reference (18)) were kept for the analysis. The 1000 most variable genes were used to compute Pearson correlation between cell lines and used to generate the heatmap (correlation scores are in Table S1).

**Whole-genome Sequencing of 4T1-4T07 cells and PDX1915-related analyses**

Genomic DNA was extracted from mammary fat pad of a female BALB/c mouse and mycoplasma-free 4T1 and 4T07 cells using the AllPrep DNA/RNA Mini Kit (Qiagen # 80204) according to the manufacturer's protocol. DNA was quantified with a Qubit Fluorometer and Qubit reagents (Thermo Fisher Scientific), and the WGS libraries were prepared using NxSeq AmpFREE kit (Lucigen). Libraries were sequenced using the Illumina NovaSeq6000 S4 v1.5 (desired sequencing depth of 100X for tumor and 30X for normal mammary fat pad samples). Raw reads were trimmed using Skewer (19) and the resulting reads were aligned to the GRCm38 mouse reference genome using BWA-MEM (20). Duplicates were marked using Picard (http://broadinstitute.github.io/picard/). Copy number alterations were identified using Sequenza (21) in each cell line sample and matched normal sample. Gains and deletions were defined using copy numbers ≥8 and ≤4, respectively. SNVs/InDels were identified by GATK's Mutect2 algorithm (22), and structural variants were called using Lumpy (23) at default parameters. Variants were annotated with mouse database using SnpEff (24). Oncoprints were constructed using the R package ComplexHeatmap (https://github.com/jokergoo/ComplexHeatmap). Variants were required to be classified as either HIGH or MODERATE impact by SnpEff to be included in the oncoprint.

To assess the expression level of *PIK3CA* in the PDX1915 in comparison with the rest of the PDXs in the same biobank, raw reads were trimmed using Trimmomatic v0.32 (25). First, the palindrome mode was used to discard adaptors and other Illumina-specific sequences from each read. Then, a four-nucleotide sliding window removes the bases once the average quality within the window falls below 30. Then, each read's first four bases were discarded. Finally, reads shorter than 30 base pairs were dropped. Cleaned reads were aligned to the combined human and mouse reference genome build mm10 and hg19 using STAR v2.3.0e (26) with default settings. Reads mapping to more than 10 locations in the genome (MAPQ < 1) were discarded. The reads that were mapped to more than 10 locations in the genome (MAPQ < 1) were removed. Towards estimating gene expression levels, primary alignments mapping to at most 2 locations (MAPQ >= 3) to exonic regions (the maximal genomic locus of each gene and its known isoforms) were quantified using featureCounts v1.4.4 (9) and the hg19 ensGene annotation set from Ensembl. Normalization (mean of ratios) of the data was performed using DESeq2 v1.14.1 (27). Multiple control metrics were obtained using FASTQC v0.11.2, samtools v0.1.20 (28), BEDtools v2.17.0 (29) and custom scripts.

**Investigating the PIK3C3 E562K mutation**

The PIK3C3 crystal structure (PDB: 3IHY) was imported to and visualized with PyMOL. The mutagenesis tool of PYMOL was used to model and visualize the (E562K) mutated protein. To investigate the effect of this mutation on mTORC1 signaling experimentally, attB sequences were first added to the *PIK3C3* sequence by performing PCR on the pcDNA4-Vps34-Flag vector (Addgene #24398). PCR product was then shuttled into the pDONR-221 vector to be subjected to site directed mutagenesis by PCR using the QuikChange II XL Site-Directed Mutagenesis Kit (Agilent Technologies, #200521) to generate the E562K mutation. The wild-type and mutated DNA sequences were shuttled to pcDNA5-pDEST-FRT-3xFlag-C-term backbone via Gateway cloning towards generating HeLa Flp-In T-REx cells expressing FLAG-tagged wild-type or mutated PIK3C3 as described before (30). Cells were seeded in 6 well-plates and expression was induced for 24 hours with tetracycline (1 μg/ml) before being processed for protein extraction and western blotting analysis, as described in the (Protein extraction, western blotting, and activity analysis) section of the materials and methods in the main text.

**Defining PIK3C3 gene essentiality (dependence) score in human breast cancer cell lines**

We accessed the DepMap portal(31) (https://depmap.org/portal) in December 2023 to define the PIK3C3 gene dependency scores in breast cancer cell lines. Scores from CAMA-1, HCC70, and T47D cell lines are shown in Figure S8G.

**Clinical datasets mining and bioinformatics analysis of clinical outcome**

To determine if the missense mutations identified in 4T1 and 4T07 cells by WGS are also present in human tumors, data from 10 PanCancer studies (32-41) (total samples = 76,639 from 75,661 patients) were accessed through the cBioportal online platfom (42,43). We specifically examined the lists of human mutations in *PIK3CA*, *RICTOR*, and *PIK3C3* for the presence of the mutations in our study.

Breast cancer gene expression and clinical data from TCGA (44) and METABRIC (45) cohorts were retrieved from the cBioPortal database. Gene set signature scores were computed using the R package GSVA(46). We selected the Hallmark_mTORC1_Signaling as a well-curated representation of mTORC1 activity-mediated signature. Each GSVA enrichment score represents the degree to which the genes in the Hallmark_mTORC1_Signaling signature are coordinately up or downregulated within a sample. For PIK3C3-related analyses, we employed a transcriptomic-based approach to generate a PIK3C3-specific gene signature. We mined two publicly available datasets (GSE206261 and GSE69747). GSE206261 (RNA-Seq) compared the transcriptome of pancreatic cancer cells treated with either DMSO or VPS34IN1 (the PIK3C3-specific inhibitor used in our study) (47). GSE69747 (Microarray) profiled lung cancer cells treated with either a PIK3C3-targeting siRNA or a scrambled siRNA(48). We reasoned that genes that are commonly downregulated when PIK3C3 activity inhibited (VPS34IN1-treated vs. DMSO-treated cells) or when PIK3C3 expression is reduced (PIK3C3-siRNA vs. scrambled siRNA) could represent a PIK3C3-specific gene signature. From these analyses, we identified 183 genes meeting this criterion. Gene Ontology term enrichment analysis of these genes showed enrichment in biological processes known to be mediated by PIK3C3(49,50), such as cell division and cytokinesis-related terms.

Breast Cancer Intrinsic Molecular Subtype (Basal, Her2, Luminal A, Luminal B) was assessed according to (Ref (51)). Heatmaps were generated using Morpheus (https://software.broadinstitute.org/morpheus). Dot plots showing differences in the Hallmark_mTORC1_Signaling scores among the molecular breast cancer subtype (Basal, Her2, Luminal A, Luminal B) were generated using GraphPad Prism Software v10.1.0 and p-values for the statistical difference between different groups were calculated using One-way ANOVA followed by Tukey multiple comparison test (p-value < 0.0001 is denoted by ****; p-value < 0.05 is denoted by * in Fig 6B and 6D).

To predict survival outcomes, the top and bottom quartile of patients were selected based on their signature score. Differences in survival were assessed using Kaplan–Meier analysis and log-rank test statistics using GraphPad Prism Software v10.1.0 and the survival and survminer R packages.

#### SUPPLEMENTARY METHODS REFERENCES


1. Li W, Koster J, Xu H, Chen CH, Xiao T, Liu JS*, et al.* Quality control, modeling, and visualization of CRISPR screens with MAGeCK-VISPR. Genome Biol **2015**;16:281
2. Hart T, Brown KR, Sircoulomb F, Rottapel R, Moffat J. Measuring error rates in genomic perturbation screens: gold standards for human functional genomics. Mol Syst Biol **2014**;10:733
3. Hart T, Moffat J. BAGEL: a computational framework for identifying essential genes from pooled library screens. BMC Bioinformatics **2016**;17:164
4. Hart T, Chandrashekhar M, Aregger M, Steinhart Z, Brown KR, MacLeod G*, et al.* High-Resolution CRISPR Screens Reveal Fitness Genes and Genotype-Specific Cancer Liabilities. Cell **2015**;163:1515-26
5. Hart T, Tong AHY, Chan K, Van Leeuwen J, Seetharaman A, Aregger M*, et al.* Evaluation and Design of Genome-Wide CRISPR/SpCas9 Knockout Screens. G3 (Bethesda) **2017**;7:2719-27
6. MacLeod G, Bozek DA, Rajakulendran N, Monteiro V, Ahmadi M, Steinhart Z*, et al.* Genome-Wide CRISPR-Cas9 Screens Expose Genetic Vulnerabilities and Mechanisms of Temozolomide Sensitivity in Glioblastoma Stem Cells. Cell Rep **2019**;27:971-86 e9
7. https://www.bioinformatics.babraham.ac.uk/projects/fastqc/.
8. Trapnell C, Roberts A, Goff L, Pertea G, Kim D, Kelley DR*, et al.* Differential gene and transcript expression analysis of RNA-seq experiments with TopHat and Cufflinks. Nat Protoc **2012**;7:562-78
9. Liao Y, Smyth GK, Shi W. featureCounts: an efficient general purpose program for assigning sequence reads to genomic features. Bioinformatics **2014**;30:923-30
10. Anders S, Huber W. Differential expression analysis for sequence count data. Genome Biol **2010**;11:R106
11. Ashburner M, Ball CA, Blake JA, Botstein D, Butler H, Cherry JM*, et al.* Gene ontology: tool for the unification of biology. The Gene Ontology Consortium. Nat Genet **2000**;25:25-9
12. Gene Ontology C. The Gene Ontology resource: enriching a GOld mine. Nucleic Acids Res **2021**;49:D325-D34
13. Mootha VK, Lindgren CM, Eriksson KF, Subramanian A, Sihag S, Lehar J*, et al.* PGC-1alpha-responsive genes involved in oxidative phosphorylation are coordinately downregulated in human diabetes. Nat Genet **2003**;34:267-73
14. Subramanian A, Tamayo P, Mootha VK, Mukherjee S, Ebert BL, Gillette MA*, et al.* Gene set enrichment analysis: a knowledge-based approach for interpreting genome-wide expression profiles. Proc Natl Acad Sci U S A **2005**;102:15545-50
15. Klijn C, Durinck S, Stawiski EW, Haverty PM, Jiang Z, Liu H*, et al.* A comprehensive transcriptional portrait of human cancer cell lines. Nat Biotechnol **2015**;33:306-12
16. Durinck S, Spellman PT, Birney E, Huber W. Mapping identifiers for the integration of genomic datasets with the R/Bioconductor package biomaRt. Nat Protoc **2009**;4:1184-91
17. Ghandi M, Huang FW, Jane-Valbuena J, Kryukov GV, Lo CC, McDonald ER, 3rd*, et al.* Next-generation characterization of the Cancer Cell Line Encyclopedia. Nature **2019**;569:503-8
